# Adjacent Z-DNA contributes to the formation of homologous DNA triplex

**DOI:** 10.64898/2026.09.29.755329

**Authors:** Mingqing Zhong, Mengqin Liu, Ran An, Yanqing Gao, Xingguo Liang

**Affiliations:** State Key Laboratory of Marine Food Processing & Safety Control, College of Food Science and Engineering, Ocean University of China, Qingdao 266404, China; Laoshan Laboratory for Marine Science and Technology, Qingdao 266235, China

**Keywords:** Z-DNA, Homologous recombination, Triplex, Circular DNA, R-loop

## Abstract

Whether the homologous parallel triplex (HP-triplex, formed by dsDNA and a homologous ssDNA) acts as an intermediate prior to DNA recombination remains controversy, owing to the intrinsic instability and lake of experimental data. Here, we report a HP-triplex that is stable at least at 35 °C under protein-free conditions. Two complementary circular ssDNAs were first hybridized at a 1:1 ratio to form an LR-chimera containing both Z-DNA and B-DNA. Then the circular homologous ssDNA (T-rich) was found to easily hybridize with this LR-chimera to form the HP-triplex (strand ratio of 2:1), involving both left- and right-handed triplex parts under the topological constraint. The third strand locates in the major groove and forms hydrogen bonds with both bases in each base pair of dsDNA without obvious strand exchange. Our data indicate that the third-strand ssDNA may first bind to the B-DNA region, followed by association with the Z-DNA region to assemble the full triplex structure. Structural simulations and analyses reveal that topological constraint imposed by the circular architecture raises the energy barrier for strand displacement, thereby stabilizing this triplex. In summary, the dynamic HP-triplex can be stabilized via a Z-B junction structure that mimics the *in vivo* one formed by negative supercoiling. These findings demonstrate that HP-triplex may act as an intermediate for DNA recombination, anti-gene strategy, and other interactions between dsDNA and homologous ssDNA or RNA.

## 1 Introduction

Whether single-stranded DNA (ssDNA) initiates strand exchange by forming a triplex with the target double-stranded DNA (dsDNA) during homologous recombination is an intriguing and challenging question^1,2^. The triplex discussed herein is designated as the homologous parallel triplex (HP-triplex). The term “parallel” indicates that the two strands with identical sequences share the same orientation^3^. In contrast, for the two classical triplexes (YR*Y and YR*R), the third strand is antiparallel to the strand with same sequence within the duplex. Specifically, a polypurine third strand aligns antiparallel to the purine strand of the duplex, whereas a polypyrimidine third strand aligns antiparallel to the pyrimidine strand. Furthermore, in classical triplexes, the third strand (which generally differs in sequence or orientation from both strands of the parent dsDNA) binds to the purine strand of the duplex via Hoogsteen hydrogen bonds^4^. By contrast, in the HP-triplex, the third strand (identical to its homologous strand in the dsDNA) forms hydrogen bonds with both bases of the Watson-Crick (W-C) base pair of the duplex DNA^5,6^. The notable feature of HP-triplex is that it does not impose so stringent sequence constraints as observed for classical triplexes, theoretically, it can form with arbitrary sequences (Figure S1, Figure 1B).

**Figure 1.**
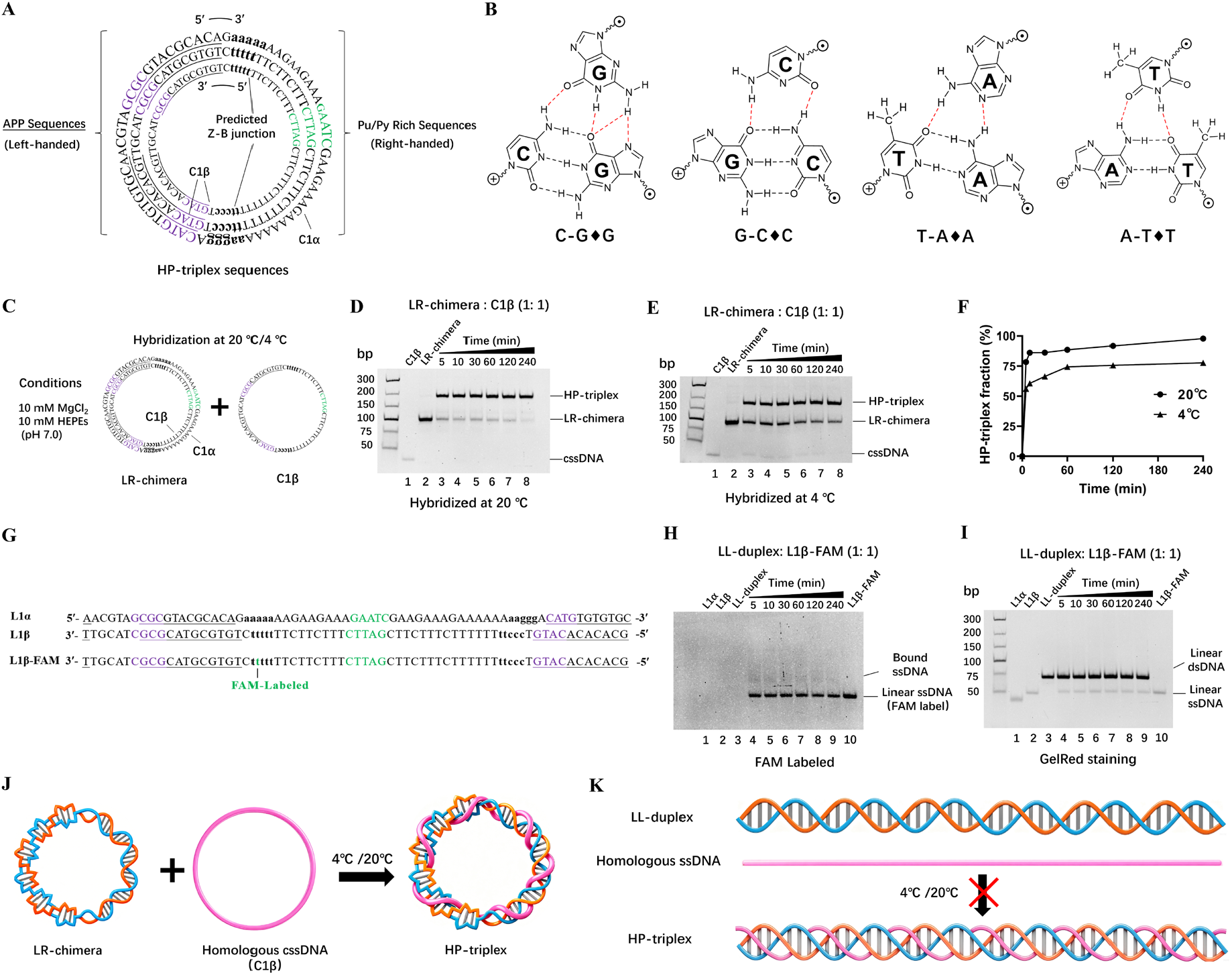
Spontaneous formation of HP-triplex from homologous cssDNA and LR-chimera. **(A)** Sequences of the cssDNAs used for HP-triplex assembly. The left 36 bp segment is designed to form left-handed DNA (approximately three turns), underline: APP sequence; The right 32 bp segment forms right-handed DNA (approximately three turns) and represents a purine-rich sequence (24 A, 6 G, 1 T, and 1 C). The bases in lowercase letters mark the predicted Z-B junction region (typically only one base pair for a Z-B junction). Hybridization of circular C1*α* (purine-rich) and C1β (pyrimidine-rich) yields the LR-chimera, and C1β serves as the homologous cssDNA. **(B)** Schematic diagrams of four types of triplons. The third base forms hydrogen bonds with both bases of the W-C base pair. Some of these hydrogen bonds are weak due to unfavorable geometry^8,25^. Black dashed lines: hydrogen bonds within the W-C base pairs; red dashed lines: hydrogen bonds formed between a base in 3^rd^ strand and the corresponding W-C base pair. ⨁ (5′-side on the paper surface) and ⊙ (3′-side on the paper surface) denote deoxyriboses. **(C)** Schematic showing the process for HP-triplex formation. After mixing LR-chimera and C1β (1:1) in the buffer (10 mM MgCl_2_, 10 mM HEPES (pH 7.0)), the mixture was incubated at 20 °C (or 4 °C) for a certain time. **(D, E)** Electrophoretic analysis at 10 °C (for incubation at 20 °C, D) and at 4 °C (for incubation at 4 °C, E). Lane 1: C1β (homologous cssDNA); Lane 2: LR-chimera; Lanes 3–8: incubation for different durations (5–240 min). Electrophoresed Gels were visualized after GelRed staining. **(F)** Time course for HP-triplex formation based on the data in (D) and (E). **(G)** Sequences of linear L1*α*, L1β, and fluorescence labeled L1β-FAM. The FAM moiety is attached to the 5-methyl group of a thymine at the B-Z junction part. **(H)** Electrophoretic analysis of linear ssDNA hybridization. Lanes 1–3: linear ssDNA of L1*α*, linear ssDNA of L1β, and their hybrid (LL-duplex), respectively; Lanes 4–9: L1β-FAM incubated with LL-duplex at 20 °C for different durations (5–240 min); Lane 10: L1β-FAM. Electrophoresis was performed at 10 °C, and the gel was imaged directly without staining. **(I)** Gel shown in panel H after GelRed staining. **(J)** Schematic illustrating HP-triplex formation between LR-chimera and homologous cssDNA. (K) Schematic showing that LL-duplex (linear duplex) fails to efficiently form triplex with homologous linear ssDNA.

HP-triplex was hypothesized as early as before 1984^5,6^ and was later termed “recombination triplex”, “parallel triplex”, or “R-DNA”^7,8^. Subsequent scattered studies demonstrated that this structure could indeed form, mostly using dsDNA shorter than 20 bp as substrates. Nevertheless, no connection to biological recombination or R-loop formation was established^1,3,9^. A. K. Shchyolkina and colleagues reported HP-triplex formation for acridine-modified 14 nt DNA sequences as detected by PAGE^10^. However, their evidence supporting HP-triplex formation was not fully accepted, because the heteroduplex dsDNA generated after strand exchange and the HP-triplex exhibited similar migration and could hardly be resolved by gel electrophoresis. HP-triplex formation mediated by recombination proteins and long ssDNA was also proposed^11,12^, yet the structural details remained unclear, and these findings could not confirm HP-triplex formation in the absence of proteins. Several rationally designed short oligonucleotides were reported to form HP-triplex via intramolecular self-folding^13,14^, but their supporting evidence, such as *T*_m_ shifts and fluorescence changes, was ambiguous. A recent piece of supportive evidence came from the group of Yan Jie (National University of Singapore, NUS), who in 2017 observed that a 572 nt ssDNA stretched by a molecular tweezer could spontaneously form HP-triplex with its homologous dsDNA^15^. Owing to the long DNA sequence used in that work, it remained difficult to determine whether partial D-loop formation occurred, rather than exclusive HP-triplex. Moreover, the artificial reduction in the entropy of ssDNA, which decreases the entropic penalty upon binding to dsDNA, may poorly reflect physiological conditions *in vivo*.

Decades of studies suggest that homologous recombination appears not to require HP-triplex formation. In this model, recombinant proteins unwind the dsDNA and slide along the duplex. The 3′ terminus of the ssDNA recognizes the complementary strand within the unwound duplex and hybridizes to form a D-loop^5,6,16–19^. Nevertheless, this homology-searching mechanism suffers from low efficiency and high energy consumption for long genomic sequences. Even within this mechanism, the unwound duplex region is typically short (3–8 nt)^20^, leaving open the possibility that HP-triplex formation assists homologous recognition. Therefore, determining whether HP-triplex can form and evaluating its stability and sequence dependence will help elucidate its role in recombination. Furthermore, homologous DNA strands are displaced during DNA replication or transcription. The displaced ssDNA or transcribed RNA may also act as the third strand to form HP-triplex again^21,22^. Elucidating the structure and formation mechanism of HP-triplex is thus critical for understanding replication and transcription, including R-loop. It is well known that many transcribed functional RNAs can serve as a homologous strand, which may exert their biological functions via HP-triplex formation. The persistent motivation driving scientists to search for HP-triplex formation arises from two aspects: first, the favorable complementarity of hydrogen bond between donors and acceptors on the third strand and duplex, rendering the structure geometrically feasible (Figure 1B); second, accumulating circumstantial evidence hinting at its existence^8,13,23–25^. Intriguingly, prior to Watson and Crick’s discovery of the DNA double helix, most scientists hypothesized that DNA adopted a triple-helical conformation^26^. Could the intuition of these eminent scientists have been entirely incorrect? We firmly believe that definitive answers to this fundamentally important scientific question can be attained through persistent efforts.

The molecular design is critically important for generating homologous triplexes. A major challenge in HP-triplex formation is that this structure generally exists as a transient intermediate which has insufficient stability and is difficult to trap. Meanwhile, selection of strand length also poses a difficulty: short strands (e.g., <30 nt) continuously undergo strand exchange and cannot be trapped in the triplex state^27,28^, whereas long strands are difficult to observe or characterize. HP-triplex may also exhibit certain sequence preferences, which have not yet been found. We noticed that the negative supercoil, which often exists in cells and readily unwinds, has not been fully considered for its effects on HP-triplex formation, especially the effect of temporarily formed Z-DNA. Herein, we propose that negative supercoiling may promote HP-triplex formation via the left-handed Z-DNA. We first hybridized two complementary 70 nt circular ssDNAs (cssDNAs) to construct a stable structure containing left-handed and right-handed helices (LR-chimera), which mimics the negatively supercoiled dsDNA under cellular physiological conditions. One extra cssDNA used to from the LR-chimera acts as the third strand to hybridize with the LR-chimera to form a circular HP-triplex. This design reduces entropic loss, introduces topological constraints that raise the activation energy for triplex dissociation. Fortunately, we obtained highly stable HP-triplexes that can be characterized by electrophoresis below 30 °C, representing substantial progress in the field of nucleic acid structural research.

## Results and Discussion

### Spontaneous HP-triplex formation between homologous circular ssDNA and circular dsDNA with a linking number (Lk) of zero

The sequence design is illustrated in Figure 1A. The starting materials consist of two fully complementary 70 nt cssDNAs, designated C1*α* and C1β (see Table S1 for the detailed sequences). The C1*α* sequence comprises two segments: the left segment is an APP-rich sequence (APP: Alternative Pyrimidine Purine) prone to Z-DNA formation, and the right segment is an A-rich sequence (24 A, 6 G, 1 T and 1 C). C1β is fully complementary to C1a. They hybridize to yields an LR-chimera with Z-DNA on one side and B-DNA on the other (L denotes left-handed, R denotes right-handed) ^29,30^. The predicted Z-B junction region is marked by lowercase bold bases. To facilitate subsequent structural analysis, three restriction enzyme sites were engineered within the sequence (color-labeled bases in Figure 1A). The synthesis of C1*α* and C1β and their hybridization to form the LR-chimera were performed according to our previously reported method (Figures S2, S3). The resulting construct was verified using specific left-handed DNA binding proteins and restriction endonuclease digestion assays, confirming the presence of Z-DNA (Figures S3, S4). Following this design, after C1β hybridizes with C1*α* to form the LR-chimera, the homologous single-stranded C1β acts as the third strand to form HP-triplex with the LR-chimera (containing left-handed and right-handed triplex segments). Notably, as shown in Figure 1B, the bases of the third strand within HP-triplex form hydrogen bonds with the bases of the other two strands, rather than forming Hoogsteen base pairs only with purine bases as observed in the widely accepted antiparallel triplex^8,25^. The triplet in HP-triplex is designated as a triplon, denoting a three-base mutually hydrogen-bonded unit, distinct from the classical triplet involving a Hoogsteen base pair.

An equimolar amount of C1β was added to the LR-chimera-containing solution with a final molar ratio of 2: 1 (C1β : C1*α*, Figure 1C). After hybridization at 20 °C for 5 min followed by electrophoretic analysis, a new, slower migrating band was observed (Lane 3). Meanwhile, the intensity of this new band gradually increased with prolonged incubation, whereas the LR-chimera band diminished markedly (Lanes 3–8). After 240 min of hybridization, almost all LR-chimera transformed to the new band (Figure 1D). These results demonstrate that single-stranded C1β can hybridize to the duplex LR-chimera, presumably forming a HP-triplex. Further increasing the C1β concentration induced no obvious change in the intensity of the new band, nor were additional new bands detected (Figure S5). This confirms that C1β and LR-chimera assemble into a 1:1 complex, namely the triplex. This hybridization reaction can also proceed at 4 °C but requires a longer incubation time. After 4 h, 20–30% of the LR-chimera remained unreacted (Figure 1E). Comparison of the time courses obtained at these two temperatures revealed that hybridization proceeded rapidly at the initial stage, followed by a slower substrate consumption rate (Figure 1F). The hybridization rate at 4 °C is slower probably because the third strand forms more stable secondary structures at lower temperature, raising the activation energy for triplex assembly.

After two linear ssDNAs of L1β and L1*α*, with identical sequences as the circular ones, were hybridized to form a linear duplex (Figure 1G), excess L1β failed to further hybridize to it (Figure S6). As ssDNAs exhibit poor binding to GelRed, they produce faint bands for all these experiments. To accurately quantify the reaction between an ssDNA and a linear duplex, the FAM-labeled ssDNA (L1β-FAM) was used as the third strand (Figure 1G). Only faint bands above the L1β-FAM band were visible prior to GelRed staining, indicating no obvious hybridization between L1β-FAM and the non-modified linear duplex (Figure 1H). After GelRed staining of the same gel (to visualize all DNA bands), the faint fluorescent band in Figure 1H overlapped with the duplex band (Figure 1I). Accordingly, no stable linear HP-triplex formed, but only tiny strand exchange might occur. Taken together, homologous ssDNA can hybridize to form the triplex only in the presence of the LR-chimera (Figure 1J), but not linear dsDNA containing exclusively right-handed helices (Figure 1K). This indicates that the left-handed segment or the topological constraint may play a critical role in the formation of this stable triplex structure.

When the circular C1*α*, bearing an A-rich sequence on its right segment, was added in excess as the third strand, no new band was observed, showing that triplex did not form between C1*α* and the LR-chimera (Figure S7). In addition, when we used LR-chimera involving a random sequence with balanced base ratios on the right side while retaining an APP-rich sequence on the left (C3*α* and C3β), no band corresponding to a triplex hybrid was detected (Figure S8). These observations indicate that HP-triplexes, if formed on a B-form dsDNA, exhibit substantial sequence selectivity. When a sequence system (C2) analogous to the C1 system was tested (the APP segment on the left of C2 forms a more stable secondary structure within the ssDNA, and the right-side sequence of C2β contains only T and C), triplex assemblies were also generated (Figure S9). This result demonstrates that HP-triplex formation is not an artifact arising from one specific sequence. Introduction of several mismatches into this sequence (replacement of the original 2 T and 2 C with A residues) still permitted triplex formation. However, the electrophoretic mobility difference between the triplex and LR-chimera decreased, and the corresponding band became more diffuse. We speculate that the mismatches reduce the structural stability of the triplex, leading to dissociation during electrophoresis (a dynamic equilibrium between strand association and dissociation) (Figure S10). This result further confirms that the stable right-handed HP-triplex may form and has sequence-specificity. Certainly, when random sequences were used on both left- and right-sides, no triplex can form. Obviously, searching for the proper sequences is the key point for further study of HP-triplex.

Notably, no obvious change in fluorescence intensity was observed upon triplex formation when staining with GelRed. This indicates that GelRed can still intercalate between the base pairs of the HP-triplex to produce strong fluorescence, although the triplex is bent to some extent due to the circular structure. It is noteworthy that dsDNA *in vivo* is usually present as a bent structure of 100–200 bp due to the stress generated by the negative supercoil. It is amazing that the triplex can form so efficiently for a circular dsDNA as short as 70 bp, and it remains stable during PAGE with normal conditions (10 mM Mg^2+^, pH 7.0, 0.1 μM of DNA). We believe the topological constraint may be the main reason to inhibit the strand displacement so that the formed HP-triplex is stable enough for detection, which is confirmed as follows.

### No strand exchange occurs between the homologous ssDNA and LR-chimera within the HP-triplex

To determine whether the third strand tightly associates with the LR-chimera duplex or undergoes strand exchange after triplex assembly, we performed restriction digestion assays on the triplex (digesting only the W-C duplex). The design of restriction sites is illustrated in Figure 2A. The HinfI site is located on the right side (B-DNA), whereas HhaI and NlaIII sites reside on the left side (Z-DNA). LR-chimera, HP-triplex, and CL-duplex (a nicked B-form circular duplex formed by a circular strand and a linear one) were individually digested (Figure 2B). After LR-chimera was digested with HinfI at 20 °C for 1 h, two new bands were generated (Lanes 5, 6, Figure 2C). The band with higher mobility was assigned to the linear duplex produced by cleavage of both strands of the LR-chimera, as its migration position matched that of the linear duplex product from HinfI digestion of CL-duplex (Lane 10). The product band migrating above the original LR-chimera was attributed to the nicked circular structure (CL-duplex) formed by cleaving only one strand in the LR-chimera, consistent with the band position of CL-duplex shown in Lane 9. This observation is fully consistent with our previous findings^30^. When HP-triplex was subjected to HinfI digestion, no new bands were produced (compare Lane 7 with Lane 8), indicating that triplex forms at least around the HinfI restriction site, where tight binding of the third strand prevents binding of the restriction endonuclease. Other two enzymes, HhaI and NlaIII (with restriction sites on the left side), could neither cleave either LR–chimera nor the HP– triplex, whereas the corresponding CL-duplex was efficiently digested (Figure 2D, E). Collectively, the LR-chimera forms a left-handed helix on the left and a right-handed helix on the right, and HP-triplex forms at least near the restriction site on the right.

**Figure 2.**
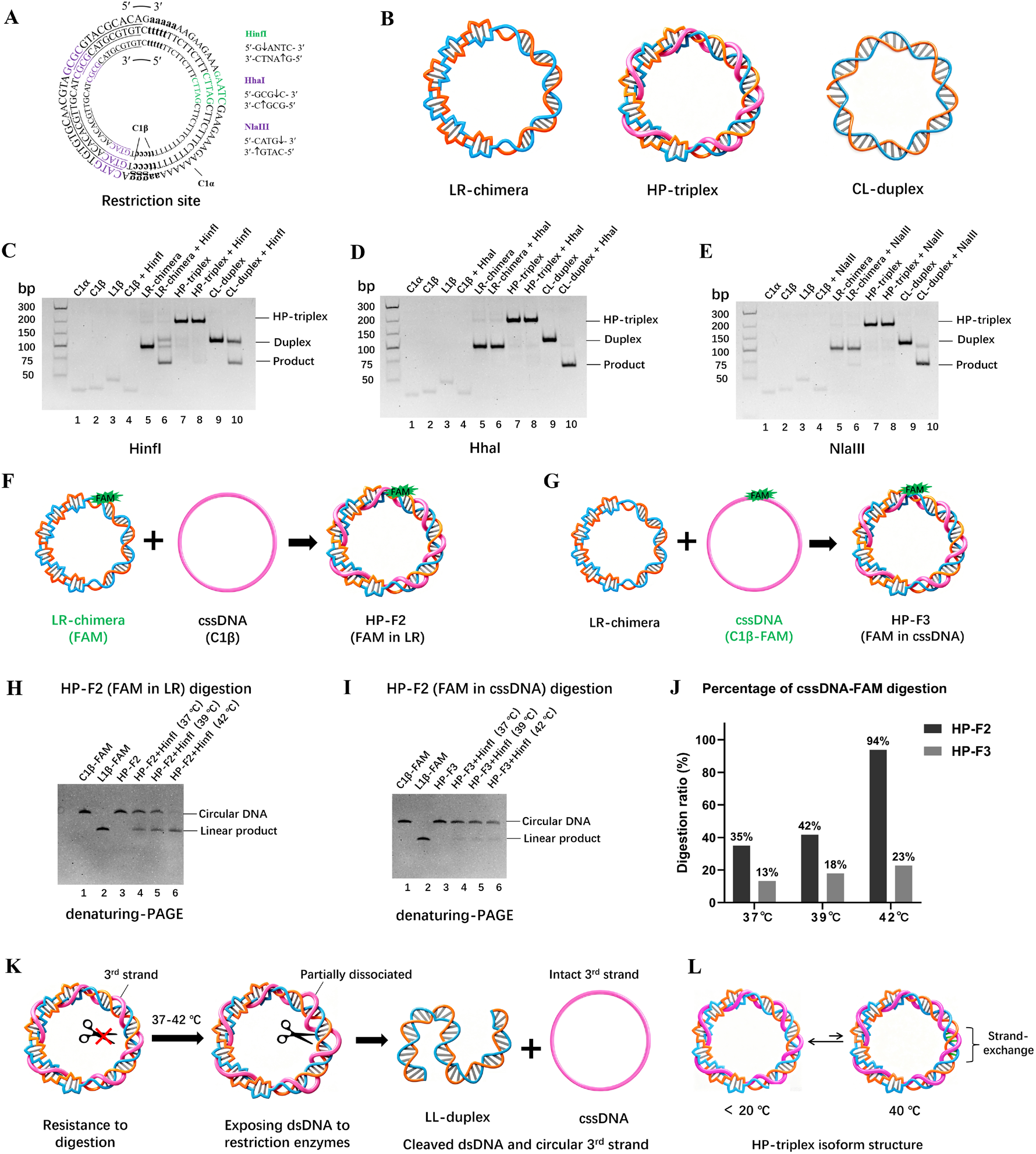
Assays of HP-triplex after digestion by Restriction endonuclease. **(A)** Schematic of restriction sites. Purple: recognition sites for HhaI and NlaIII (Z-DNA region); green: HinfI recognition site (B-DNA region). **(B)** Schematic diagrams of LR-chimera, HP-triplex, and CL-duplex (nicked circular duplex). **(C–E)** Digestion results for HinfI, HhaI, and NlaIII, respectively. Lanes 1–3: C1*α*, C1β, and L1β, respectively; Lane 4: C1β mixed with restriction enzyme; Lanes 5, 7, 9: LR-chimera, HP-triplex, and CL-duplex, respectively; Lanes 6, 8, 10: After digestion of samples shown in lanes 5, 7 and 9, respectively. Conditions: 0.1 μM DNA, 0.5 U/μL restriction enzyme, 20 °C, 1 h; 10% PAGE (10 mM MgCl_2_, pH 7.5, run at 10 °C), visualized after GelRed staining. **(F, G)** Schematic diagrams of HP-triplex assembly with C1β-FAM incorporated as 2^nd^ strand (HP-F2) or 3^rd^ one (HP-F3). **(H, I)** HinfI digestion at various temperatures. Lane 1: C1β-FAM; Lane 2: L1β-FAM; Lane 3: HP-triplex; Lanes 4–6: digestion at 37 °C, 39 °C, and 42 °C for 12 h, respectively. Analyzed on 15% dPAGE (8 M urea, run at 20**–**25 °C), imaged without staining. **(J)** Quantification of C1β-FAM digestion efficiency from panels H and I. **(K)** Proposed digestion mechanism of HP-triplex at 37**–**42 °C. Partial dissociation of 3^rd^ strand exposes the duplex region, which is then cleaved to yield the linear duplex. **(L)** Schematic of dynamic strand exchange within the HP-triplex.

Considering a DNA triplex usually exhibits lower thermal stability than a duplex, and only the third strand dissociates at a relatively higher temperature, we performed restriction digestion at elevated temperatures (37 °C, 39 °C, and 42 °C) to identify whether the third strand exchanged with its homologous one (Figures 2F–2J). It is supposed that even partial dissociation will expose the dsDNA for the digestion. To differentiate the second strand and the third one with the same sequence, we used FAM-labeled circular ssDNA (C1β-FAM) as either the second (HP-F2, Figure 2F) or the third one (HP-F3, Figure 2G) for triplex assembly prior to digestion. As expected, at elevated temperatures, HinfI digestion was observed after 12 h, and the digestion efficiency increased with temperature (Lanes 4–6, Figure S11). Interestingly, when C1β-FAM served as the second strand, the obtained digestion product (linear duplex) exhibited strong fluorescence without staining, confirming that C1β-FAM remained in duplex as the second strand (Lanes 4–6, Figure S11A). When C1β-FAM was used as the third strand during triplex assembly, the digestion product showed almost no FAM fluorescence, whereas the circular ssDNA released after cleavage displayed intense fluorescence, demonstrating that C1β-FAM retained its identity as the third strand during digestion (Lanes 4–6, Figure S11B). To better compare the digestion profiles of C1β-FAM, a denaturing-PAGE was performed (Figure 2H–J). The results clearly showed that the digestion efficiency reached 35–94% when C1β-FAM acted as the second strand, markedly higher than the value of 13–23% observed for C1β-FAM as the third strand (Lanes 4–6, Figure 2H–J). Collectively, these results indicate that even at elevated temperatures, the second strand remains predominantly in W-C base pairing, and little or no strand displacement occurs for the third strand.

Based on the above data, we propose a model for the high-temperature restriction digestion process of the triplex, as illustrated in Figure 2K. At elevated temperatures above 37 °C, the third strand of the HP-triplex dissociates, or partially dissociates, from the duplex to expose the duplex region. As shown in Figure 1, linear duplexes hardly bind to the third single-stranded. Therefore, once the exposed duplex is cleaved into a linear form, the circular third strand is released. On one hand, strand exchange is barely detectable for assembled HP-triplex at low temperatures below 20 °C. On the other hand, a small amount of strand exchange may occur upon incubation at 37–42 °C (Figure 2L). Such strand exchange may involve only a portion of the 70 nt sequence and exists in a dynamic equilibrium between strand invasion and reversion. Since the *T*_m_ of the LR-chimera is generally greater than 60 °C^29,30^, and the structure is additionally wrapped by the third strand, the probability of LR-chimera duplex’s melting at 37–42 °C is extremely low.

When electrophoresis was performed at 40 °C without digestion, strong fluorescence retained in the LR-chimera band (C1β-FAM as the second strand) after HP-triplex melting during electrophoresis, and the dissociated the third strand exhibited almost no fluorescence (Lanes 4–6, Figure S12A). In the case where C1β-FAM acted as the third strand, intense fluorescence was observed at the position corresponding to circular ssDNA, whereas no fluorescence was detected at the position corresponding to LR-chimera (Lanes 4–6, Figure S12B). These observations are consistent with the digestion results, demonstrating that even at 40 °C, the third strand does not undergo strand exchange. Instead, only triplex melting occurs, whereby the third strand dissociates from the LR-chimera. It is noteworthy that the concentration becomes much lower during electrophoresis due to the diffusion at both loading and running steps.

### CD and AFM characterization of the HP-triplex structure

The CD spectra of LR-chimera was measured at 20 °C (Figure 3A). Compared with the CL-duplex (right-handed construct with identical sequences), LR-chimera exhibited obvious shifts and alterations in CD peaks. After subtracting the B-DNA signal from the spectrum of LR-chimera, the CD signal corresponding to left-handed lh-DNA was obtained, showing characteristic negative peak at 295 nm and positive peak at 267 nm (the positive peak of B-DNA appears at 280 nm) (Figure 3B), similar as we reported previously^30^. CD spectral changes of LR-chimera were monitored over the temperature range of 10**–**80 °C. No obvious spectral variation was observed for LR-chimera from 10 to 60 °C, whereas marked changes occurred at 70 °C and 80 °C. These data indicate that LR-chimera remains stable below 60 °C (Figure 3C).

**Figure 3.**
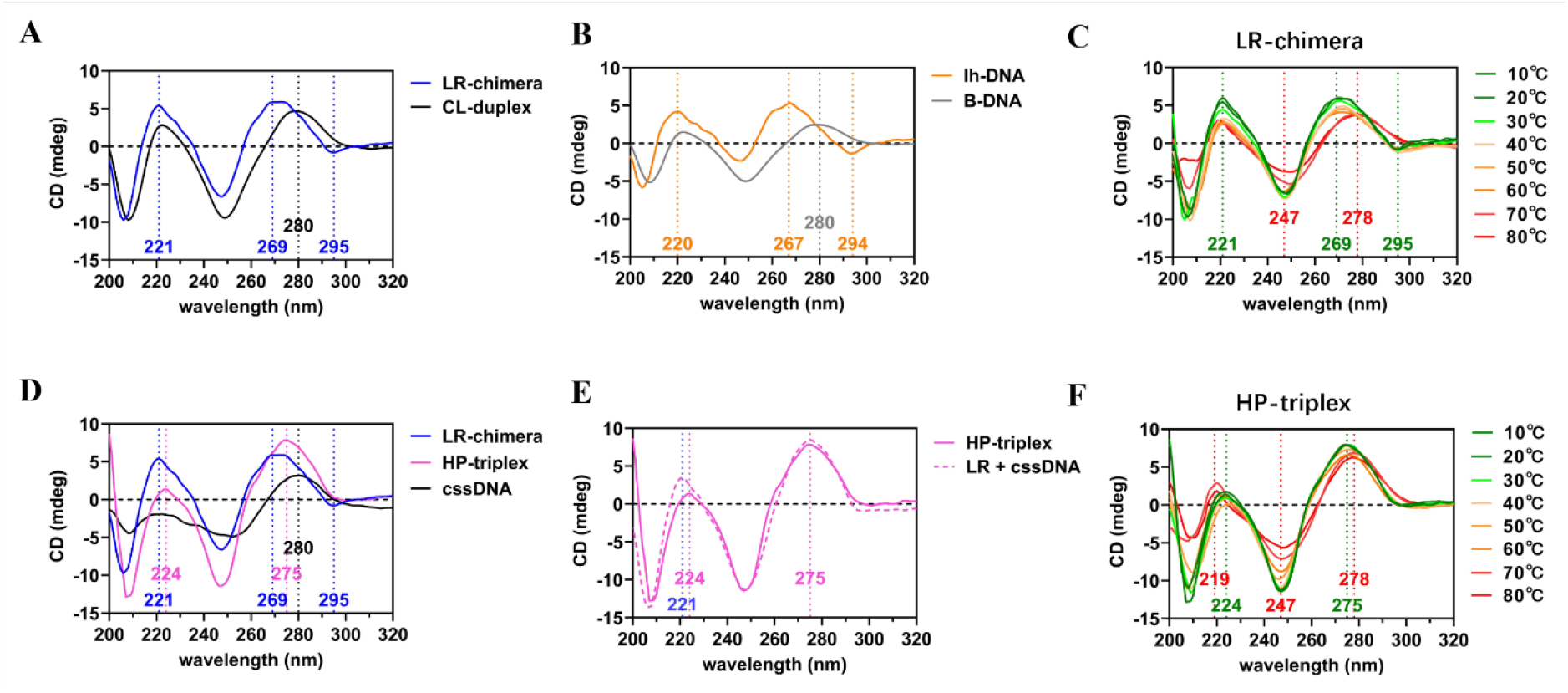
CD spectra of LR-chimera and HP-triplex. **(A)** Blue: LR-chimera; Black: CL-duplex (nicked circular duplex). Measured at 20 °C. **(B)** Orange: lh-DNA (LR – 0.47 × CL), CD signal of LR-chimera after subtraction of the B-DNA contribution; Gray: B-DNA (0.53 × CL), CD signal of B-DNA with the same length as lh-DNA. **(C)** CD spectra of LR-chimera recorded at various temperatures (10–80 °C). **(D)** Blue: LR-chimera; Pink: HP-triplex; Black: cssDNA (C1β). Measured at 20 °C. **(E)** Solid pink line: HP-triplex; Dotted pink line: CD signal of LR-chimera plus cssDNA. **(F)** CD spectra of HP-triplex recorded at different temperatures (10–80 °C). Conditions: 4 μM DNA, 10 mM MgCl_2_, 10 mM HEPES (pH 7.0).

Although the CD spectra of the HP-triplex differed markedly from those of the LR-chimera and cssDNA at 20 °C (Figure 3D), direct summation of the CD signals of LR-chimera and cssDNA yielded a spectrum that nearly overlapped with that of HP-triplex (Figure 3E). This observation suggests that the duplex and single-stranded components undergo no substantial structural rearrangement upon triplex formation. No obvious spectral variation was detected from 10 to 40 °C, and only minor alterations were observed upon heating to 50 °C and 60 °C (Figure 3F). These subtle changes may be attributed to dissociation of the third strand above 50 °C. As expected, the CD signal decreased substantially at 70 °C and 80 °C, corresponding to melting of the LR-chimera scaffold.

AFM was also employed to characterize the molecular structures of LR-chimera and HP-triplex (Figure S13). LR-chimera adopted a roughly circular morphology with a height of ~1.2 nm, which was slightly lower than the theoretical value of 2 nm. This discrepancy likely arose from measurement artifacts under dry-state imaging conditions^31^. Compared with LR-chimera, HP-triplex exhibited a better-defined ring shape and an increased height of ~1.6 nm. This finding matched the expectation that wrapping of homologous cssDNA around LR-chimera thickens the helical assembly. Collectively, the CD and AFM results support the conclusion that HP-triplex is formed by hybridization between the homogenous cssDNA and the LR-chimera.

### HP-triplex remains stable at least at 35 °C

To characterize the thermal stability of HP-triplex, high resolution melting (HRM) assays were performed to monitor fluorescence changes upon temperature elevation^29,30,32,33^. The underlying principle is that the small molecule dye EvaGreen binds and emits fluorescence only when DNA secondary structures form, but ssDNA without a strong secondary structure shows negligible fluorescence because it barely interacts with EvaGreen. The results show that LR-chimera binds EvaGreen and generates strong fluorescence at lower temperatures, and gradually decreases as the duplex melts at higher temperatures (Figure 4A). A sharp fluorescence drop was observed at approximately 60–70°C, and the obtained *T*_m_ is 66.6 °C (Figure 4A). HRM data for HP-triplex reveal that the fluorescence intensity is markedly lower for HP-triplex than for LR-chimera (less than half), indicating that the binding of the third strand hinders EvaGreen’s binding to the major groove of the dsDNA. As temperature increases, conversely, fluorescence rises slowly over the range of 35–63 °C. This trend probably deviates from the dissociation of the third strand so that more EvaGreen can bind. Similar to LR-chimera, fluorescence decreases rapidly above 63 °C, and a nearly identical *T*_m_ of 66.8 °C is obtained. Clearly, these results also support that HP-triplex can form, and it is stable at least below 35 °C. This is also confirmed by the following UV melting experiments.

**Figure 4.**
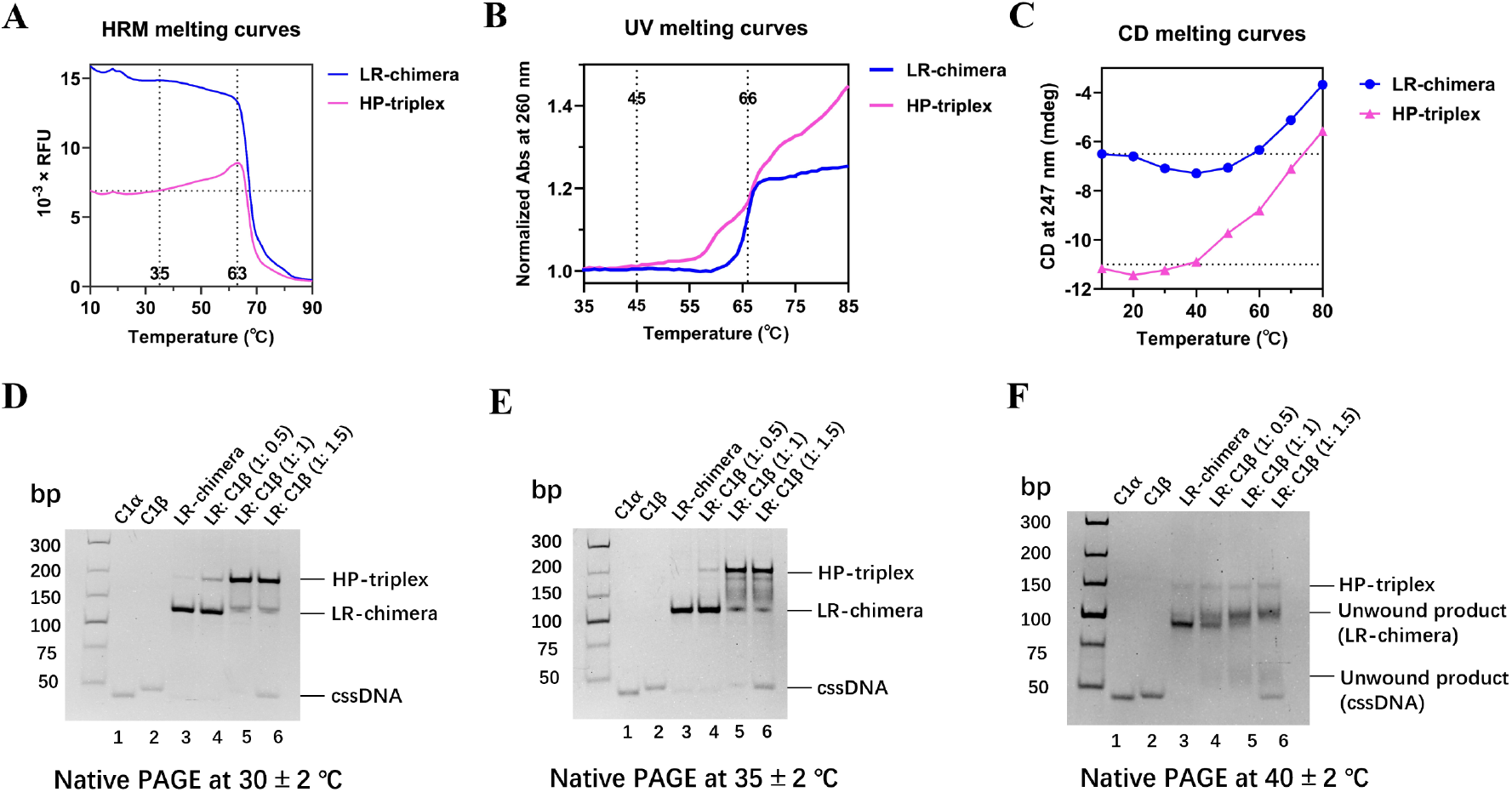
Characterization of the thermal stability of HP-triplex. **(A-C)** HRM, UV and CD melting curves, respectively. Blue: LR-chimera; Pink: HP-triplex. Conditions: pH 7.0, 1 μM (A) or 0.5 μM (B) HP-triplex. Data for (C) are from Figures 3C and 3F at 247 nm. **(D-F)** Electrophoresis at 30 °C, 35 °C, and 40 °C, respectively. Lanes 1-3:C1*α*, C1β, and LR-chimera, respectively;Lanes 4-6:Hybridization of LR-chimera with 0.5, 1.0, and 1.5 folds of C1β. Conditions: 0.1 μM LR-chimera, 10 mM MgCl_2_, 10 mM HEPES (pH 7.0); 10% native-PAGE (10 mM MgCl_2_, pH 7.5), Stained by GelRed.

The UV melting curve of HP-triplex exhibits two distinct transitions (Figure 4B). The first transition occurs from 45 to 66 °C, corresponding to dissociation of the third strand, and the second transition spans 60 to 70 °C. This profile matches the canonical melting features of triplex structures ^13,34,35^. Likewise, analysis of temperature-dependent CD signals at 247 nm identifies signal changes for HP-triplex between 40 and 60 °C that are not observed for LR-chimera, demonstrating that the dissociation of the HP-triplex third strand primarily occurs between 40 and 60 °C (Figure 4C). This observation is consistent with the restriction digestion results, where HP-triplex is susceptible to cleavage at temperatures above 37 °C (Figure 2).

The stability of HP-triplex at varying temperatures was further investigated by controlling the gel temperature during native-PAGE using circulating water. HP-triplex remains nearly undissociated at electrophoresis temperatures of 10 °C, 20 °C and 30 °C (Figure 4D and Figure S14). Only faint smearing bands can be observed between the bands corresponding to HP-triplex and LR-chimera at 35 °C (Lanes 4 – 6, Figure 4E). At 40 °C, HP-triplex dissociates almost completely into LR-chimera and single-stranded products during electrophoresis (Lanes 4–6, Figure 4F). These results demonstrate that HP-triplex is sufficiently stable below 30 °C, and begins dissociation at 35 °C under electrophoresis. It is noteworthy that once the third strand dissociates, it migrates away from the duplex in the electric field due to its distinct mobility. In addition, the pH in the gel is 7.5, which also destabilizes the triplex to some extent. Consequently, the thermal stability of HP-triplex measured by native-PAGE is substantially lower than its real stability. HinfI digestion assays also revealed that the cleavage rate of HP-triplex was much slower than that of LR-chimera even at 42 °C (Figure 3). This indicates that the third strand remains predominantly bound to LR-chimera at 42 °C. Therefore, the *T*_m_ of HP-triplex is estimated to be at least above 42 °C.

### Topological constraint from the circular third strand stabilizes the HP-triplex

To verify whether the topology of homologous cssDNA is required for HP-triplex formation, linear homologous ssDNA (L1β) was incubated with LR-chimera instead of the circular one (Figure 5A). After incubation of L1β and LR-chimera at 20 °C for 4 h, the band corresponding to LR-chimera could not observed. On the contrary, a new band with slightly reduced mobility (matching that of CL-duplex) and another band migrating identically to C1β were observed (Lanes 5–7, Figure 5B). This result demonstrates that the linear third strand displaced the circular ssDNA of the second strand within LR-chimera (Lanes 4–8). Furthermore, the displaced homologous circular ssDNA failed to form a triplex with the newly formed CL-duplex (B-DNA). This indicates that stable triplex structures can only form when all the three strands are circular, and the topological constraint is necessary. HP-triplex cannot form regardless of whether the linear ssDNA acts as the second or the third strand.

**Figure 5.**
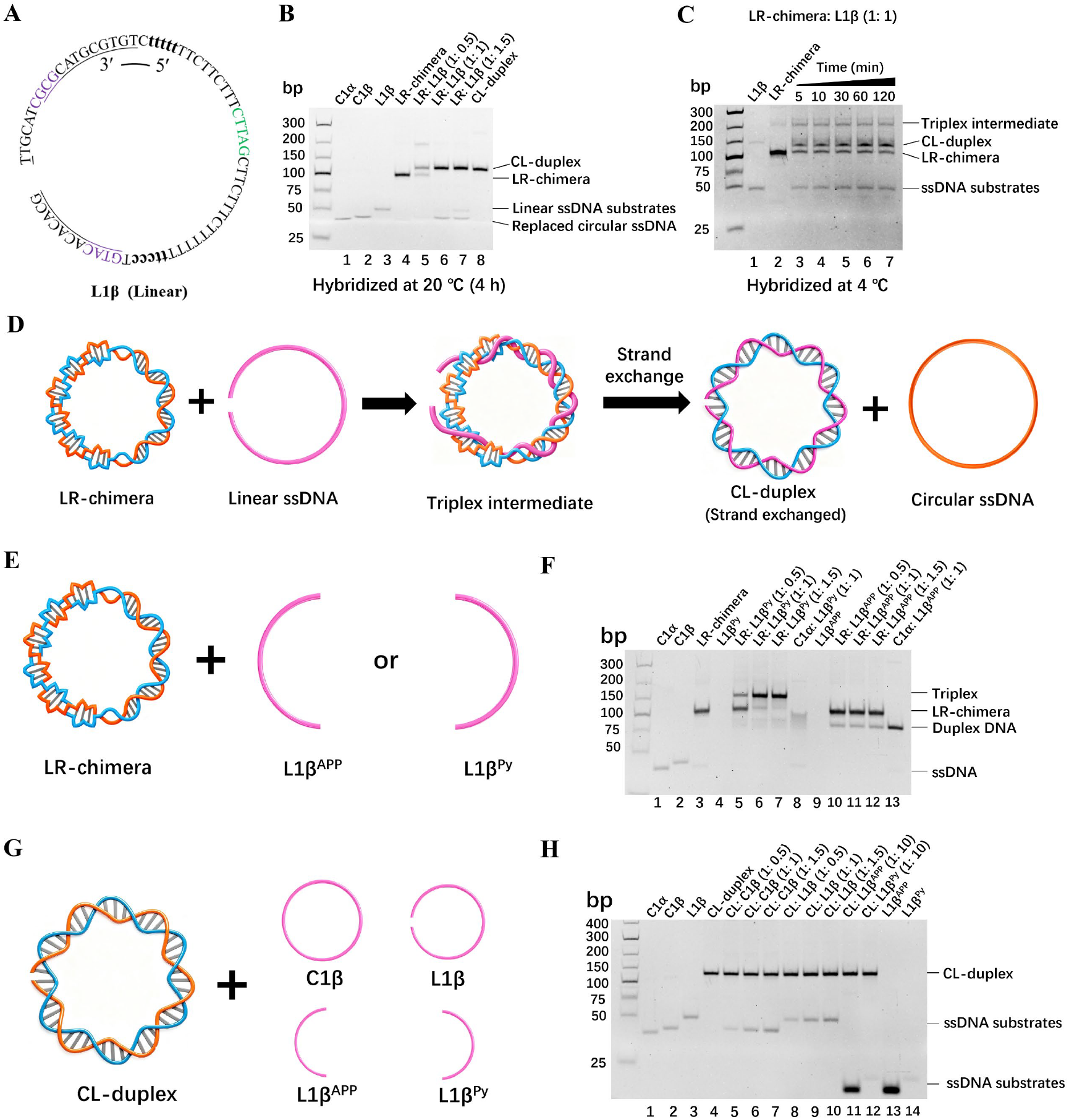
Mechanism underlying HP-triplex formation. **(A)** Schematic of linear ssDNA (L1β). **(B)** Hybridization of LR-chimera with L1β. Lanes 1–3: C1*α*, C1β, and L1β; Lane 4: LR-chimera; Lanes 5–7: LR-chimera mixed with L1β at various molar ratios; Lane 8: CL-duplex. **(C)** Hybridization of LR-chimera with L1β at 4 °C. Lane 1: L1β; Lane 2: LR-chimera; Lanes 3–7: LR-chimera incubated with L1β (1:1) for different durations at 4 °C. Electrophoresis performed at 4 °C. **(D)** Scheme for strand exchange between linear ssDNA and LR-chimera. After LR-chimera hybridizes with linear ssDNA to form a triplex intermediate, strand displacement occurs to produce CL-duplex and circular ssDNA. **(E)** Scheme for hybridization of LR-chimera with L1β^Py^ (right segment) or L1β^APP^ (left segment). **(F)** Results corresponding to panel E. Lanes 1-2: C1*α*, C1β; Lane 3: LR-chimera; Lane 4: L1β^Py^; Lanes 5–7: LR-chimera mixed with L1β^Py^ at various molar ratios; Lane 8: L1β^Py^ mixed with C1*α* (1:1); Lane 9: L1β^APP^; Lanes 10–12: LR-chimera mixed with L1β^APP^ at various molar ratios; Lane 13: L1β^APP^ mixed with C1*α*. **(G)** Scheme for mixing CL-duplex with C1β, L1β, L1β^Py^, or L1β^APP^. **(H)** Results corresponding to panel G. Lanes 1–2: C1*α*, C1β; Lane 3: L1β; Lane 4: CL-duplex; Lanes 5–7: CL-duplex mixed with C1β at various molar ratios; Lanes 8–10: CL-duplex mixed with L1β at molar various ratios; Lane 11: CL-duplex mixed with L1β^APP^ (10-fold concentration); Lane 12: CL-duplex mixed with L1β^Py^ (10-fold concentration); Lane 13: L1β^APP^; Lane 14: L1β^Py^. Other conditions (except as otherwise noted): 0.1 μM DNA, 10 mM MgCl_2_, 10 mM HEPES (pH 7.0), incubation at 20 °C for 4 h, electrophoresis at 10 °C, pH7.5.

Studies on the rate of strand displacement by the linear third strand revealed that the homologous cssDNA within LR-chimera was displaced over 80% within 5 min at 20 °C, indicating very rapid dynamics (Lane 3, Figure S15). A band was observed far above the LR-chimera band with substantially lower mobility than CL-duplex, and it faded over time and eventually disappeared completely (Lanes 3–7). This species was likely a triplex byproduct formed during the strand displacement, because the displaced circular C1β could hybridize with the remaining unreacted LR-chimera to generate the HP-triplex. Once all the strand displacement is finished, almost no LR-chimera remained (all C1*α* hybridized with L1β), and HP-triplex almost disappeared completely (Lane 7, Figure S15). To better resolve this strand-exchange process, the reaction temperature was lowered to 4 °C to slow strand-exchange kinetics (Figure 5C). A distinct HP-triplex band was observed. This confirms that both LR-chimera and C1β were present throughout the displacement process, and they formed HP-triplex gradually (Lanes 3–7, Figure 5C).

Can linear L1β form HP-triplex or not? To address this question, FAM-labeled linear ssDNA (L1β-FAM) was used for further investigation (Figure S16). At 4 °C of hybridization, bands corresponding to the triplex and the duplex both showed the fluorescence signal from L1β-FAM. This demonstrates that at least a fraction of L1β can form a triplex with LR-chimera, which remains relatively stable at low temperature (Lanes 3–7, Figure S16). These results imply that strand displacement proceeds via an intermediate triplex structure prior to full strand exchange. Significantly, when LR-chimera bearing random sequences was used (random sequences cannot form HP-triplex), no strand exchange was observed during incubation with linear L3*α* or L3β at 20 °C (Figure S17). This indicates that strand exchange between homologous linear ssDNA and LR-chimera at low temperatures requires the formation of sufficiently stable HP-triplex (Figure 5D). Again, the strand-exchange process is hard to occur through a mechanism that the duplex melts first so that the ssDNA can hybridize,^18,19^ because LR-chimera is very stable at 4 °C. These results and analysis provide experimental evidences supporting HP-triplex as an intermediate during recombination.

### The structure of B-DNA linked to Z-DNA contributes greatly to stabilize the HP triplex

To investigate the contribution of the Z-DNA moiety to HP-triplex assembly, short linear oligonucleotides corresponding to the Z-side segment (L1β^APP^) or B-side segment (L1β^Py^) of the third strand was incubated with LR-chimera (Figure 5E). After hybridization between L1β^Py^ and LR-chimera at 20 °C for 4 h, a new band was observed, suggesting triplex formation within the right-handed helical region (Lanes 5–7, Figure 5F). Comparison with the hybrid between L1β^Py^ and C1*α* (Lane 8) demonstrates that L1β^Py^ did not displace the homologous circular C1β completely from the LR-chimera to form a duplex. In contrast, no band with reduced mobility was observed when L1β^APP^ was used, indicating that L1β^APP^ fails to bind the left-handed region of LR-chimera (Lanes 10–12). It is well known that ssDNA (without other stable secondary structures) adopts highly flexible right-handed conformations. Accordingly linear ssDNA is difficult to associate with left-handed duplex to form a triplex without topological constraint. These findings also suggest that during assembly of the full circular HP-triplex from homologous circular ssDNA and LR-chimera, triplex formation initiates on the right-helical side before propagation into the complete HP-triplex. What roles does the Z-DNA segment play in HP-triplex assembly and stabilization?

When CL-duplex (an all-right-handed B-DNA formed by a circular ssDNA and its linear complementary strand) was used in place of LR-chimera and incubated with circular or linear homologous ssDNA (Figure 5G), no binding was observed for either substrate (Lanes 5–10, Figure 5H). No band shifts were detected even when the ssDNA concentration was increased to 10-fold relative to CL-duplex (Lanes 11, 12). These results indicate that L1β^Py^, which carries only the right-side sequence, cannot form stable HP-triplex with its homologous B-DNA. Although transient binding can occur (entropically unfavorable), the complex preferentially dissociates (entropically favorable). Stable HP-triplex assembly between the B-DNA segment of LR-chimera and homologous ssDNA (circular or linear, including L1β^Py^) requires the presence of Z-DNA directly linked to the B-DNA region. Furthermore, strand displacement is suppressed when the third strand is circular.

Accordingly, a plausible mechanism is proposed for the formation of circular HP-triplex (Figure 6A). The circular third strand first associates with the right-handed segment of LR-chimera to generate a triplex intermediate. Its remaining portion then wraps leftward around LR-chimera to yield the full HP-triplex. After HP-triplex formation, topological constraints act such that even partial strand displacement occurs, the displaced strand is confined within the minor groove and cannot immediately dissociate (Figure 6A). The system therefore prefers to revert to its original state, i.e., HP-triplex. Whether comparable partial strand displacement takes place within the left-handed segment remains undetermined (Figure 6B). Even upon complete displacement at moderately elevated temperatures such as 30 °C, the released circular ssDNA can reassociate with LR-chimera to form new HP-triplex, establishing a dynamic equilibrium. Consequently, HP-triplex appears highly stable at least at 30 °C.

**Figure 6.**
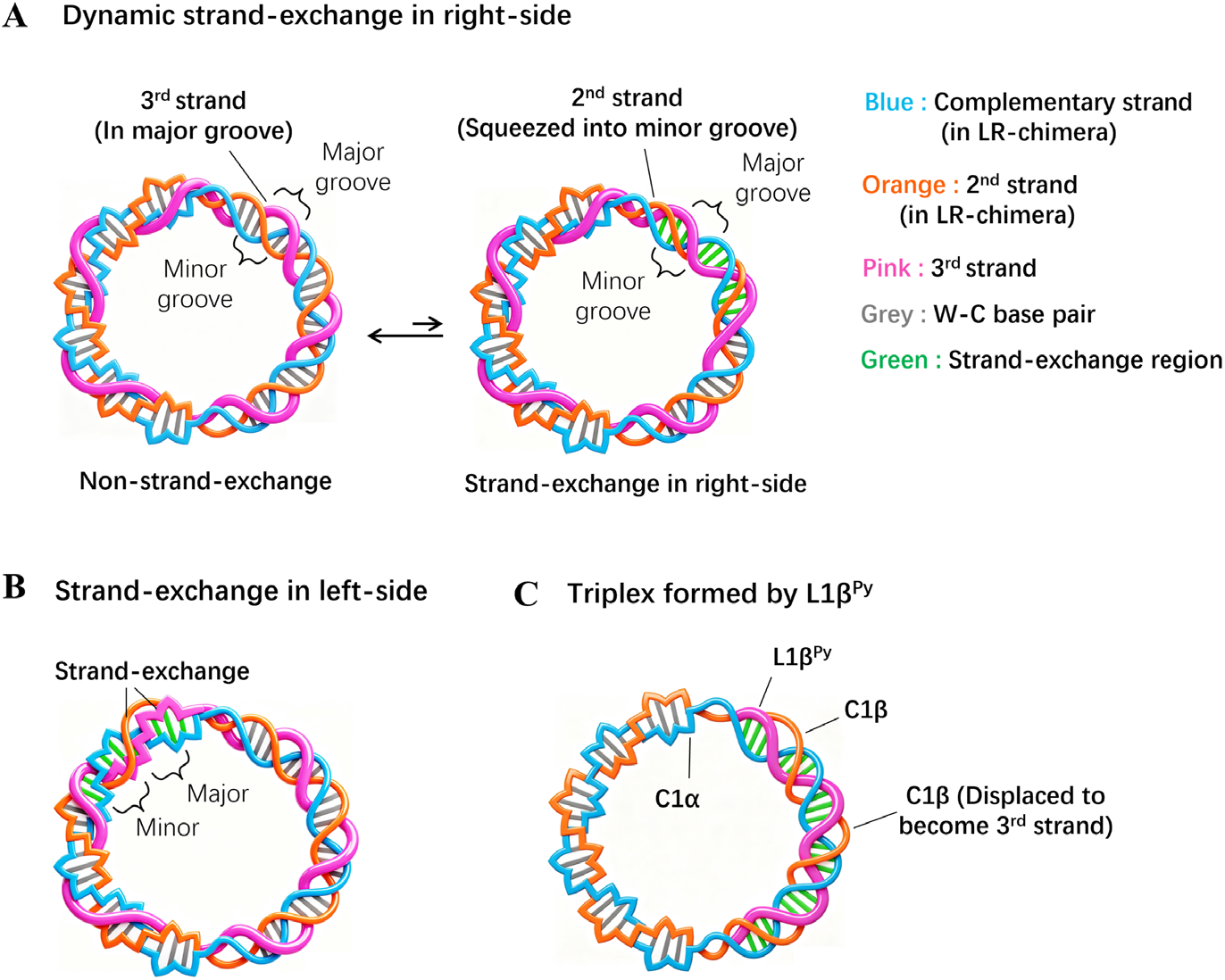
Mechanism for HP-triplex stabilization under topological constraints. (A) Schematic illustration for dynamic strand exchange of partial sequences on the right-handed side of HP-triplex. Left: HP-triplex without strand exchange; Right: partial strand exchange of 3^rd^ strand within the right-handed helical segment. (B) Schematic showing dynamic strand exchange of partial sequences on the left-handed side. (C) Schematic of the complex formed between L1β^Py^ (pink) and LR-chimera. The right-handed helical segment of the C1β strand (orange) is displaced by L1β^Py^ as 3^rd^ strand. L1β^Py^ forms a W-C paired duplex with C1*α* at the right-handed region. In all panels, gray bars denote hydrogen bonds of W-C base pairs within LR-chimera; green bars denote hydrogen bonds of W-C base pairs formed after strand exchange.

In contrast, linear L1β^Py^ can hybridize to the right-handed region of LR-chimera, as explained as follows. HP-triplex forms first, followed by strand invasion, yielding a B-form right-handed helix with the circular C1*α*. The displaced C1β does not dissociate promptly because the Z-DNA segment remains sufficiently stable. Meanwhile, the right-side segment of displaced C1β acts as the third strand, wrapping on major groove to form HP-triplex after leaving the minor groove (Figure 6C). A cooperation effect exists between the left-side Z-DNA and formation of the right-side HP-triplex, preventing simultaneous dissociation of whole strand. Obviously, if dissociation occurs on only one side, the structure tends to revert back to its original conformation which is more stable.

Do hydrogen bonds depicted in Figure 1B also form within the left-handed region of the HP-triplex? To address this question, another third-strand of C2β-m^APP^ was designed. It contains 20 bp mismatches evenly distributed along its left segment relative to the first strand C2*α*, while its right segment remains fully complementary (Figure S18A). Evidently, C2β-m^APP^ cannot form hydrogen bonds with the Z-DNA duplex region of LR-chimera and exhibits nearly no complementarity. Nevertheless, after hybridization of C2β-m^APP^ with LR-chimera, a distinct triplex band was observed via electrophoresis (electrophoresis performed at 10 °C) (Lanes 6 – 8, Figure S18B). This demonstrates that complementarity of the left-handed segment is not so essential for triplex formation. Instead, topological constraints likely play a more critical role. As described above, topological constraints suppress strand displacement, thereby stabilizing the circular HP-triplex. This finding further confirms that the right-side sequence of the third strand possesses an intrinsic strong capacity to form HP-triplex at least at low temperatures. In the absence of topological constraints, however, substantial entropic penalties raise the free energy of the triplex structure, and the complex tends to dissociate.

Notably, at a higher temperature of 27 °C, electrophoretic analysis of these mismatch-containing triplexes revealed that the mismatch-bearing system produced a faster-migrating band below the triplex band. This observation indicates that this system undergoes melting more readily and gradually releases LR-chimera compared with the fully matched system (Lanes 5, 6, Figure S18C). This result demonstrates that binding between the homologous ssDNA strand with Z-DNA also occurs to form HP-triplex at the left part by forming hydrogen bonds (triplons) as illustrated in Figure 1B. Because the major groove of Z-DNA protrudes outward from the helix, binding of the third-strand there encounters less steric hindrance than binding to B-DNA. It is noteworthy that the left segment consists mainly of APP or random sequences (not T-rich). However, it remains unclear whether Z-DNA facilitates HP-triplex formation on B-DNA.

Binding of the circular HP-triplex to a Z-DNA-binding protein (the Zα binding domain) was also examined (Figure S19 and Figure S4B). Weak binding (smearing bands) was detected only at high protein concentrations (> 10 μM). Although HP-triplex binds the protein more readily than B-DNA, we propose that HP-triplex is poorly recognized by Z-DNA-binding proteins. This is consistent with the fact that the Zα domain primarily interacts with the major groove, which is occupied by the third strand in HP-triplex^36–38^. In summary, the Z-DNA segment plays dual roles in HP-triplex formation, including providing topological constraints to suppress strand displacement, as well as stabilizing the HP-triplex by forming triplons.

Studies investigating the effect of magnesium ion concentration on the formation of the circular HP-triplex were also performed (Figure S20). In the presence of 1 mM Mg^2+^, a distinct new band was still observed above the LR-chimera band, although it migrated very close to the LR-chimera band. This indicates that HP-triplex can still form even at a Mg^2+^concentration as low as 1 mM. Analytical ultracentrifugation (AUC) results also confirmed the formation of HP-triplex at 1 mM Mg^2+^ (data not shown). The lower mobility on PAGE at higher Mg^2+^concentration may be caused by neutralizing the negative charge on the triple helix. Obviously, higher density of negative charge on triplex can cause stronger interaction between phosphate and Mg^2+^ than that on DNA duplex. At lower magnesium concentrations, this effect becomes much weaker so that the mobility becomes close to that the duplex. It is possible that some magnesium ions may bind tightly to the HP-triplex at elevated Mg^2+^ concentrations (e.g. 10 mM). An alternative explanation invokes a rapid equilibrium between triplex dissociation and association during electrophoresis. A fraction of the triplex population exists as duplex and single-stranded species throughout electrophoresis, leading to faster migration. Since the rate of triplex formation and dissociation in this dynamic equilibrium is much higher than the electrophoretic migration rate, the band is not so smearing. Anyway, since it has a different mobility from that of dsDNA, the HP triplex still formed at 1 mM Mg^2+^.

At 5 mM Mg^2+^, the HP-triplex band migrated identically to that at 10 mM, showing that HP-triplex is stable as the ion-strength close to physiological conditions. Notably, this topologically constrained HP-triplex forms across a broad pH range of pH 6.0–8.0 (Figure S21), indicating that formation of a classical antiparallel triplex is impossible in this case. It is well known that antiparallel triplexes containing C within the triplex-forming oligonucleotide (TFO) exhibit strong pH dependence. Such structures are particularly difficult to form at a pH above 7.0^39,40^. In addition, the presence of A and G impairs the ability of pyrimidine-rich TFO to form classical triplexes with dsDNA. The stability of the HP-triplex at low Mg^2+^ concentrations (1–5 mM) and elevated pH further confirmed that it is a homogenous parallel triplex. It is noteworthy that the inversion of the third-strand (C1β) to generate an antiparallel triplex with LR-chimera would introduce more than ten mismatched base pairs, which is extremely unstable.

### Structures of the right-handed HP-triplex and the left-handed one

The 12 bp B-DNA-side sequence of C1*α* and 14 bp APP sequence (approximately one turn of triplex helix) were selected. Structural simulations of right-handed and left-handed HP-triplex were separately performed (Figure 7). During structural modelling, the W-C duplex was kept fixed, and the third strand was initialized with the same conformation of its homologous strand within the duplex. This third strand was introduced into the major groove of the duplex, and the three bases of each triplon were constrained to co-plane. After addition of TIP3P explicit water and 10 mM magnesium ions, molecular dynamics simulations were carried out under the CHARMM force field.

**Figure 7.**
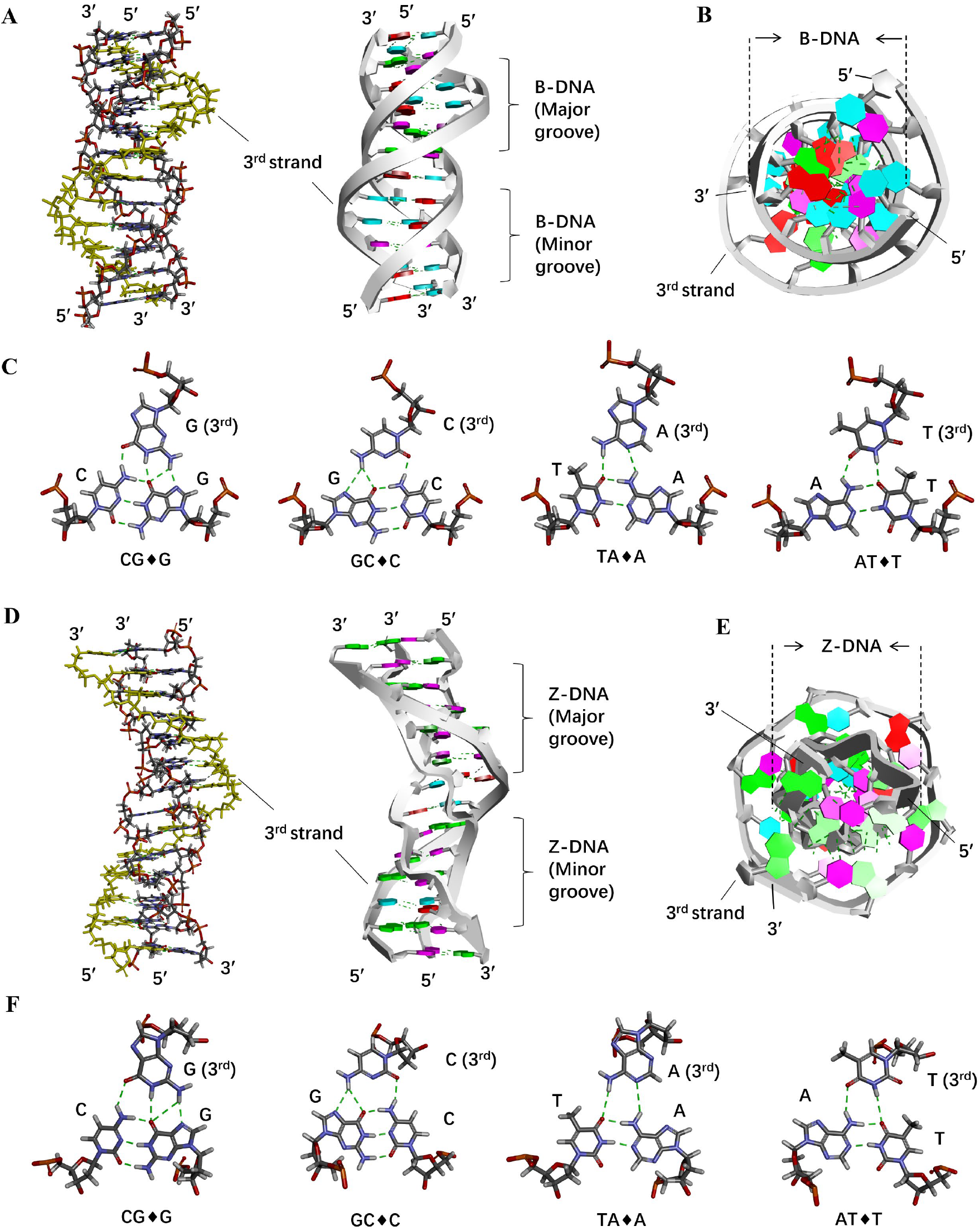
Molecular simulation structures of right-handed and left-handed HP-triplex. **(A)** Structural model of the right-handed HP-triplex. Left: stick view; Right: ribbon view. The homologous third strand has the sequence 5′-TCTTCGATTCTT-3′. 3^rd^ strand resides in the major groove of the duplex. Bases are colored differentially: red, A; light blue, T; purple, C; green, G. **(B)** Top view of panel A. **(C)** Schematic of the four triplons that constitute HP-triplex. Hydrogen bonds between bases are shown as green dashed lines. **(D)** Structural model of the left-handed HP-triplex. Left: stick view; Right: ribbon view. The homologous third strand has the sequence 5′-CGTGCGTACGCGCG-3′. 3^rd^ strand resides in the major groove of the left-handed helix. Base coloring is identical to panel A. **(E)** Top view of panel D. **(F)** Schematic showing the four triplons constituting the left-handed HP-triplex. Hydrogen bonds are shown as green dashed lines.

The simulation results reveal that the third strand binding to the major groove of the duplex in the right-handed HP-triplex. The phosphate backbone of the third strand protrudes outward from the major groove, yielding a triplex with an obvious larger diameter than the duplex (Figures 7A, 7B). The bases of the third strand simultaneously interact with both base pairs of the W-C duplex through hydrogen bonds. The four types of triplons are shown in Figure 7C (largely identical to those depicted in Figure 1B). Notably, all glycosidic linkages connecting the third-strand bases to deoxyribose adopt the *anti* conformation. Substantial size differences between purines and pyrimidines may introduce steric hindrance upon third-strand binding. Accordingly, for the HP-triplex with a right-handed conformation involving purine-rich (and pyrimidine-rich) sequences, the phosphate backbone at purine positions is pushed outward to a greater extent than that at pyrimidine positions, probably reducing the triplex stability of a mixed sequence involving both purines and pyrimidines (Figure S22).

Compared with other triplons, the hydrogen bond (N6–H…N1) formed between TA♦A shows notable angular distortion^8,25^. The lone-pair electrons angle of the acceptor N1 is approximately 90°, while the ideal angle is 180° (Table S2). This may reduce the stability of the TA♦A triplon and weaken base stacking between adjacent base pairs. This can explain to some extent the sequence selectivity of HP-triplex formation that the A-rich sequences cannot act as the third strand to form HP-triplex. Although the hydrogen bonding of the AT♦T triplon is also imperfect, T-rich β strands readily serve as the third strand for HP-triplex formation. A plausible explanation is that the methyl group of thymine on the third strand engages in hydrophobic interactions within the major groove, mitigating the entropic penalty accompanying HP-triplex formation. Further investigations are required to fully elucidate the mechanism underlying sequence-dependent stability of HP-triplex.

Surprisingly, the structure of the left-handed HP-triplex was easily obtained by simulations without special settings, and base-pairing patterns closely resembled those of the right-handed HP-triplex (Figure 7D, 7E). Compared with the right-handed HP-triplex (Figures 7A, 7B), the phosphate backbone of the third strand protrudes further outward into the major groove, corresponding to a slightly larger separation between the backbone of the third strand and those of the other two strands. This feature likely arises from the shallow major groove of Z-DNA. Furthermore, within the left-handed HP-triplex, purines in the third strand adopt the *syn* glycosidic conformation, while pyrimidines adopt the *anti* conformation, consistent with that observed in Z-DNA (Figure 7F). Notably, the left-handed HP-triplex forms a Z-like structure with every two base pairs as a repeating unit. Nevertheless, the backbone of the third strand is relatively smooth and lacks the zig-zag conformation characteristic of Z-DNA (Figure 7D). This structural feature suggests that the third strand is relatively flexible and relaxed, with no prominent steric hindrance.

All bases in the third strand (TFO) adopt the *anti* conformation in classical antiparallel triplexes. This may explain why the third strand cannot contain both purines and pyrimidines. The mixed bases would introduce steric hindrance to weaken the base pairing or induce abnormal phosphate backbone geometry (Figure S22). On the other hand, when a 6 bp non-APP sequence is inserted into the middle of the left-handed HP-triplex, simulated structures reveal that glycosidic bonds of adjacent nucleosides in the non-APP region also follow alternating *syn–anti* conformations, and the duplex retains the zig-zag structure (Figure S23). Since pyrimidines are energetically disfavored in the *syn* conformation, Z-DNA containing non-APP sequences exhibits reduced stability. Analogously, the third strand of the left-handed HP-triplex adopts alternating *syn–anti* conformations, which lowers HP-triplex stability upon incorporation of non-APP sequences. Unexpectedly, the overall architecture of the simulated left-handed HP-triplex is insensitive to sequence composition, indicating that left-handed HP-triplex can potentially form with a wide range of sequences, albeit with variable stability.

As shown in Figures 1 and 4, stable HP-triplex was obtained by introducing topological constraints and Z-DNA. Previous reports have shown that recombination-associated proteins such as RecA can bind Z-DNA, though the underlying mechanism remains unclear^41–43^. From our viewpoint, following nucleosome disassembly, a stretch of 10–20 bp Z-DNA may form^44^, and this Z-DNA segment enables the recognition of the dsDNA by ssDNA or ssRNA. The formed HP-triplex can be stabilized with the assistance of relevant proteins^38^. Perhaps at sites where Z-DNA is formed, or in adjacent B-DNA regions, RecA or other corresponding proteins facilitates more effectively the recognition of homologous dsDNA by ssDNA and promotes the subsequent strand exchange reaction.

In the absence of proteins, HP-triplex formation and dissociation persist in a dynamic equilibrium. Typically, only partial regions within an ssDNA sequence readily assemble into stable HP-triplex, while the remaining portion imposes topological constraints that suppress strand displacement. In the presence of recombination proteins, proteins can bind ssDNA (with bases generally exposed) and form HP-triplex with dsDNA. Partial protein dissociation then initiates strand displacement. Upon HP-triplex formation under the action of relevant proteins, strand displacement proceeds immediately. This accounts for the general difficulty in capturing such structures by crystallography or other approaches.

It is well known that the ssDNA and RNA strands within cells readily form or unwind secondary structures, and proteins facilitate the unwinding of such secondary structures to enable recombination. Studies have shown that before ssDNA participates in homologous strand displacement, recombination proteins extend ssDNA to approximately 1.5 times the length of canonical B-DNA^9,15,45^. Our results demonstrate that the third strand of HP-triplex winds within the major groove of the duplex and widens the helix (Figure S13 and Figure 7). The phosphate backbone lengths of the simulated models were quantified. The third strand measures approximately 84 Å (10 bp), around 1.28-fold longer than that for the duplex (approximately 66 Å) (Table S3). This indicates that elongation of the homologous ssDNA is also required for HP-triplex formation, consistent with previous reports^15^. It suggests that the elongation of ssDNA prior to homologous strand exchange may function to properly align the ssDNA with its homologous duplex, thereby facilitating base recognition and pairing, i.e., HP-triplex assembly. The third strand of the Z-form triplex also undergoes elongation, though less pronounced than observed for the B-form triplex.

Some reports imply that recombination occurs more frequently at certain special sequences *in vivo*. For instance, D-loops and R-loops (structures that remain undissociated after HP-triplex strand displacement) are enriched at CpG island promoters, transcription termination sites, and immunoglobulin class-switch regions. These structures participate in transcriptional regulation, chromatin remodeling, and DNA damage signaling^46^. The 3′ single-stranded TTAGGG repeats of telomeric DNA invade the homologous duplex to form T-loops, which block telomerase-mediated telomere elongation and mediate telomere maintenance mechanisms^47^. Telomeric RNA TERRA also forms R-loops to engage in telomere maintenance^48^.These findings are consistent with our observation that HP-triplex formation exhibits pronounced sequence preference. This preference may underpin the markedly elevated frequency of homologous strand exchange at these genomic regions *in vivo*. Investigating the sequence specificity governing HP-triplex formation may help explain the cellular distribution of D-loops and R-loops and predict sites where recombination takes place.

## Conclusion

Under topological constraints, HP-triplexes can persist stably under physiological ionic conditions, and left-handed HP-triplex conformations are achievable. The circular HP-triplex contains both left- and right-handed helical segments, in which the third strand binds in the major groove of the duplex and recognizes double-stranded DNA via hydrogen bonding. This study demonstrates for the first time that HP-triplexes can form and be stabilized by suppressing strand displacement, offering a strategy to overcome the difficulty of capturing this transient intermediate state. Strand displacement is inhibited because topological constraints tether the two termini of partially displaced ssDNA, hindering its release with a negligible entropy gain. Based on this principle, strand displacement remains unfavorable for long dsDNA substrates even when HP-triplexes form. Nevertheless, ssDNA readily forms stable secondary structures that substantially raise the activation energy for HP-triplex assembly, which may partially account for the historical difficulty in obtaining HP-triplex species.

Negative supercoiling *in vivo* promotes the formation of localized Z-DNA. Such Z-DNA regions may facilitate HP-triplex formation and favor local strand displacement. This process becomes more efficient in the presence of relevant proteins. The proteins stretch ssDNA, disrupting its secondary structure and enabling improved recognition and pairing with the duplex DNA. Subsequent recombination is accomplished through a zipping mechanism. We propose that HP-triplexes represent important dynamic structures mediating and regulating genetic functions in living organisms, particularly for the recognition of dsDNA by single-stranded nucleic acids including RNA. For instance, the formation of R-loops and the recognition and regulation of dsDNA by functional RNAs such as lncRNAs may be associated with HP-triplex assembly. Notably, this work only provides initial evidence that HP-triplexes can stably exist in the absence of proteins. Further investigations are required to characterize their properties and biological functions. Although the 70 nt circular DNA duplexes and triplexes in this system exhibit high curvature, they partially recapitulate the bending features of cellular dsDNA. It is anticipated that insights from this study will stimulate rapid advances in the field of nucleic acid triple helices in the near future.

## Materials and methods

### Materials

All oligonucleotides used in this study were purchased from Jinweizhi Biotechnology Co., Ltd. (Suzhou, China) or Sangon Biotech (Shanghai, China), T4 polynucleotide kinase and exonuclease I from Abclonal Technology (Wuhan, China), T4 DNA ligase and ATP from Thermo Scientific (Pittsburgh, PA, USA), *HinfI, HhaI*, and *NlaIII* from New England Biolabs Inc. (Ipswich, MA, USA), Ultra GelRed from Vazyme (Nanjing, China), and EvaGreen dye from Biotium, Inc. (Fremont, CA, USA). All other chemicals were purchased from Sigma–Aldrich (St. Louis, MO, USA) or Sinopharm Chemical Reagent Co., Ltd. (Shanghai, China).

### Preparation and purification of circular DNA

A phosphate group was added to the 5′ end of the linear DNA as follows. The 20 μL reaction mixture was made and contained 40 μM linear DNA, 1× T4 polynucleotide kinase buffer A (70 mM Tris-HCl, 10 mM MgCl_2_, 5 mM DTT, pH 7.6 at 25 °C), 250 μM ATP, and 0.5 U/μL T4 polynucleotide kinase. After incubated at 37 °C for 12 h, the kinase was inactivated by heating at 65 °C for 20 min.

For the self-circularization (without a splint), the 20 μL reaction mixture was prepared containing 10 μM linear DNA with 5’ phosphate and 0.2× T4 DNA ligase buffer (8 mM Tris-HCl, 2 mM MgCl_2_, 2 mM DTT, 100 μM ATP, pH 7.8 at 25 °C). The mixture was incubated at 80 °C for 3 min, then cooled to 20 °C at a rate of 0.1 °C/s and incubated at 20 °C for 5 min. Subsequently, 1 μL of T4 DNA ligase (5 U/μL) was added, and the mixture was incubated at 37 °C for 24 h. The ligase was inactivated by heating at 65 °C for 10 min.

For the ligation using a splint, the 20 μL reaction mixture was made containing 2 μM 5’-phosphated linear DNA, 4 μM splint, and 0.25× T4 DNA ligase buffer (10 mM Tris-HCl, 2.5 mM MgCl_2_, 2.5 mM DTT, 125 μM ATP, pH 7.8 at 25 °C). After mixed, the solution was incubated at 80 °C for 3 min, then inserted into an ice bath for 3 min. After 1 μL of T4 DNA ligase (5 U/μL) was added, the mixture was incubated at 37 °C for 24 h. The ligase was inactivated by incubation at 65 °C for 10 min.

Subsequently, 1 μL exonuclease I (10 U/μL) and 1 μL 10× exonuclease I buffer (670 mM glycine-KOH, 67 mM MgCl_2_, 10 mM DTT, pH 9.5 at 25 °C) were added to 20 μL circularized DNA solution. After mixed, the mixture was incubated at 37 °C for 24 h to remove linear by-products. The exonuclease I was inactivated by incubation at 80°C for 20 min. An equal volume of extraction reagent (phenol-chloroform-isoamyl alcohol, 25:24:1, v/v/v) was added to the digested products to remove residual proteins. The DNA was precipitated with ethanol to eliminate ions, resuspended in ultrapure water, and stored at −20 °C for subsequent experiments.

### HP-triplex formed by hybridization of circular ssDNA and LR-chimera

For the LR-chimera, the 10 μL solution was prepared containing 0.5 μM C1*α*, 0.5 μM C1β, 10 mM HEPES buffer (pH 7.0 at 25 °C), and 10 mM MgCl_2_. The mixture was incubated at 80 °C for 5 min, then gradually cooled to 50 °C at a rate of 0.1 °C /s and incubated at 50 °C for 15 min. Subsequently, the temperature was further decreased to 20 °C at a rate of 0.1 °C /s and maintained at 20 °C for 10 min. Other duplex DNAs were prepared using the identical procedure.

For the HP-triplex assembly, LR-chimera and C1β were mixed at a 1:1 molar ratio in 10 mM HEPES buffer (pH 7.0 at 25 °C) containing 10 mM MgCl_2_. The mixture was incubated at 20 °C or 4 °C for 5–240 min according to the experimental design. At each time point, the samples were immediately loaded onto a 10% native-PAGE gel (supplemented with 10 mM MgCl_2_, pH 7.5) for electrophoresis to separate single-stranded and double-stranded DNA. Other triplex-forming sequences were assembled via the same procedure.

### Restriction enzyme digestion assay

According to the method described above, 9 μL of double-stranded or triple-stranded DNA (0.2 μM) was prepared. Subsequently, 0.5 μL of restriction enzyme (10 U/μL) and 0.5 μL of CutSmart Buffer (500 mM potassium acetate, 200 mM Tris-acetate, 10 mM magnesium acetate, 1000 µg/mL recombinant albumin, pH 7.9 at 25 °C) were added. After mixing, the sample was incubated at 25 °C for 1 h, or at 37–42 °C for 12 h according to the experimental design. Then, the digested products were loaded onto either a 10% native-PAGE gel (supplemented with 10 mM MgCl_2_, pH 7.5) or a 15% denaturing-PAGE gel (supplemented with 8 M urea). Electrophoresis of the native-PAGE gel was run at 10 °C, whereas the denaturing-PAGE was run at 20–25 °C. After staining with GelRed (FAM direct imaging), the gel was imaged using a Bio-Rad Gel Doc™ XR+ imaging system (USA).

### Electrophoretic mobility shift assay with Zα and Z-22 (specific Z-DNA binding protein)

According to the methods described above, LR-chimera, CL-duplex, or HP-triplex was prepared separately. Subsequently, 10 μL of each DNA sample (0.1 μM) was mixed with either Zα protein (concentrations ranging from 0.1 to 20 μM) or Z-22 protein (concentrations ranging from 0.05 to 0.5 μM) in binding buffer (10 mM HEPES, 10 mM MgCl_2_, pH 7.0 at 20 °C) and incubated at 20 °C for 2 h. The mixtures were then loaded onto a 10% native PAGE gel (containing 10 mM MgCl_2_) and subjected to electrophoresis at 10 °C. After GelRed staining, the gels were imaged using a Bio-Rad Gel Doc™ XR+ imaging system (USA).

### Circular dichroism (CD) Spectroscopy

CD spectroscopy measurements were performed on a JASCO-1500 spectropolarimeter (Japan). According to the method described above, 300 μL of double-stranded or triple-stranded DNA (4 μM) were prepared. CD spectra were recorded using a quartz cuvette with a 1 mm optical path length, scanning a wavelength range from 200 to 320 nm at a scanning speed of 100 nm/min. The measurements were conducted at temperatures ranging from 10 °C to 80 °C according to the experimental design. The buffer without DNA was used as a blank control to subtract background signals. Each sample was scanned in triplicate, and the average spectrum was calculated.

### Melting analysis by UV

According to the method described above, 100 μL of LR-chimera and HP-triplex (0.5 μM) were prepared. UV melting curves of DNA were measured at 260 nm using a UV-1800 spectrophotometer (Shimadzu, Japan) with a temperature controller (Shimadzu, Japan), while the temperature was ramped from 30 °C to 85 °C at a rate of 1 °C /min. The absorbance of the DNA was recorded using a micro quartz cuvette with a 1 cm optical path length. Every measurement was performed three times and the average value was calculated. All absorbance data were normalized to the absorbance at 30 °C. Melting curves were plotted with temperature as the abscissa and normalized absorbance as the ordinate.

### High resolution melting (HRM) analysis

HRM curves were measured using a Piko Real-Time PCR system (Thermo Scientific, Finland). According to the method described above, 10 μL of double-stranded or triple-stranded complexes (1 μM) were prepared. Subsequently, 0.5 μL of EvaGreen (20×) nucleic acid dye was added and the mixture was mixed thoroughly. The samples were then placed into a real-time fluorescence quantitative PCR instrument for melting curve detection. The changes of fluorescence intensity were recorded at 0.3 °C increment while samples were heated from 10 °C to 95 °C at a rate of 1 °C/min. The excitation and emission wavelengths of EvaGreen were 500 nm and 530 nm, respectively. The melting temperature (*T*_m_) values of the LR-chimera were calculated from the the first derivative of the HRM melting curves.

### Atomic force microscope (AFM)

AFM imaging was performed using an SPM-NanoA microscope (Shimadzu, Japan). LR-chimera and HP-triplex samples were prepared at a concentration of 10 nM in 10 mM HEPES buffer (pH 7.0) containing 10 mM MgCl_2_. Subsequently, 10 μL of the sample solution was dropped onto a freshly cleaved mica surface incubated for 1 min. The surface was rinsed three times with deionized water to remove residual unbound samples. After drying with nitrogen gas, AFM scanning was conducted to characterize the nucleic acid structures.

### Analytical Ultracentrifugation (AUC)

Analytical ultracentrifugation with fluorescence detection was used to analyze the samples. The experiments were performed on an XPN-100 ultracentrifuge (Beckman Coulter) equipped with a fluorescence detector (PW-FDS001) developed by Beijing Protein Win Technology Co., Ltd. According to the method described above, FAM-labeled 400 μL HP-triplex samples (0.1 μM) were prepared. An An50Ti rotor and 12 mm centerpieces were used for sample detection. The experimental rotor speed was set to 45,000 rpm, and the data acquisition time interval was set to 126 s. The fluorescence detector parameters were set as follows: the detector gain was 0.7 V, and the laser output power was 20 mW. During the experiments, changes in the fluorescence signal of the samples during ultracentrifugation were recorded in real time, and the corresponding sedimentation curves were obtained. The processing and analysis of the experimental data were completed based on AUCAgent (https://www.frcbs.tsinghua.edu.cn/download/software/133).

### Establishment of theoretical triplex model

The theoretical models of right-handed and left-handed HP-triplexes were constructed and visually inspected using BIOVIA Discovery Studio Visualizer (Dassault Systèmes BIOVIA, San Diego, USA). Standard all-atom molecular dynamics simulations were performed using AMBER with the CHARMM force field. Briefly, the B-DNA duplex (5′-TCTTCGATTCTT-3′) and Z-DNA duplex (5′-CGTGCGTACGCGCG-3′) were first built according to the given sequences. The homologous third strand was then inserted into the major groove of each duplex to assemble the initial triple-helical structures, and atomic coordinates were exported as PDB files for subsequent simulation preparation^49^. The structures were solvated in a TIP3P explicit water box with a minimum buffer distance of 7 Å from the nucleic acid atoms to the box boundary. MgCl_2_ (10 mM) was added to reproduce the experimental ionic conditions. During model construction, positional restraints were applied to the duplex backbone to maintain double-helix conformation. Dihedral and harmonic restraints were imposed on triplon bases to preserve their coplanar arrangement. After preliminary optimization, a multi-stage relaxation strategy was adopted. The initial restraints were gradually reduced to a final force constant lower than 3 kcal·mol^−1^·Å^−2^ to allow natural structural relaxation. Subsequently, the system temperature was gradually elevated from 50 K to 300 K over a 10 ps heating phase. The SHAKE algorithm was applied to constrain all hydrogen-containing bonds, enabling a stable 2 fs time step. Finally, sequential NVT and NPT equilibrations were performed to achieve full system equilibration prior to production molecular dynamics simulations.

## Acknowledgements

We acknowledge the assistance of the Analysis and Testing Center, College of Food Science and Engineering, Ocean University of China. We thank the Core Facility for Biomolecule Preparation and Characterization at the Technology Center for Protein Sciences, Tsinghua University, for assistance with analytical ultracentrifugation fluorescence data analysis. We also gratefully acknowledge Beijing Protein Win Technology Co., Ltd. for providing the analytical ultracentrifugation fluorescence detector used in this study.

## Author Contributions

Xingguo Liang proposed the key idea. Mingqing Zhong and Xingguo Liang conceived and designed the research and drafted the manuscript. Mingqing Zhong performed the experiments. Other authors discussed the findings and assisted with manuscript revision. All authors reviewed the manuscript and approved the final version.

## Supplementary Information

Figures S1 to S23 Tables S1 to S3

