## Supplementary Information for "Adjacent Z-DNA contributes to the formation of homologous DNA triplex"

**Table S1.** Sequences of ssDNA and splint used in this study.

| Name | Sequences (5′ → 3′) * | Length (nt) |
|---|---|---|
| **C1α** | AACGTAGCGCGTACGCACAGAAAAAAAGAAGAAAGAATCGAAG AAAGAAAAAAAAGGGACATGTGTGTGC | 70 |
| **C1β** | GCACACACATGTCCCTTTTTTTTCTTTCTTCGATTCTTTCTTCTTTT TTTCTGTGCGTACGCGCTACGTT | 70 |
| **C2α** | GTACGCACGGATAAAAAGAAGAAAGAAAAGAAGAAAGAAAAAA ATGGGACACGTACGCACAACCAAGTGC | 70 |
| **C2β** | GCACTTGGTTGTGCGTACGTGTCCCTTTTTTTTCTTTCTTCTTTTCT TTCTTCTTTTTTTCCGTGCGTAC | 70 |
| **C3α** | GCGTGCGTGCGGTCTAATACGACTCACTATAAATCGAACATCAGC AAACGGACACCTCGCATTCAACTTAGAAT | 74 |
| **C3β** | ATTCTAAGTTGAATGCGAGGTGTCCGTTTGCTGATGTTCGATTTAT AGTGAGTCGTATTAGACCGCACGCACGC | 74 |
| **C2β-m$^{Py}$** | GCACTTGGTTGTGCGTACGTGTCCCTTTTTTTTCTTTCTTAATTTCT TTCTTAATTTTTTCCGTGCGTAC | 70 |
| **C2β-m$^{APP}$** | ACGATTAATTACGTATCAGGATACATTTTTTTTCTTTCTTCTTTTCTT TCTTCTTTTTTCCAGTACGGGC | 70 |
| **L1α-sp** | TACGTTGCACAC | 12 |
| **L1β-sp** | GTGTGCAACGTA | 12 |
| **C2β-m$^{APP}$-sp** | AATCGTGCCCGT | 12 |
| **L1β$^{Py}$** | TTTTTTTCTTTCTTCGATTCTTTCTTCTTTTT | 32 |
| **L1β$^{APP}$** | TCTGTGCGTACGCGCTACGTTGCACACACATGTCCC | 36 |

*The underlined section indicates the APP (alternating purine-pyrimidine) sequence.

**Table S2.** Angles and distances of triplon hydrogen bonds in right-handed and left-handed helices.

| Right helix | | | | | Left helix | | | | |
|---|---|---|---|---|---|---|---|---|---|
| **Triplons** | **H-bonds** | **DHA (°)[a]** | **HAY (°)[b]** | **Distance (Å)** | **Triplons** | **H-bonds** | **DHA (°)** | **HAY (°)** | **Distance (Å)** |
| **CG♦G** | N4-H···O6 | 162.80 | 113.88 | 1.83 | **CG♦G** | N4-H···O6 | 135.84 | 129.70 | 2.35 |
| | N1-H···O6 | 163.56 | 164.95 | 1.89 | | N1-H···O6 | 171.47 | 150.14 | 2.15 |
| | N2-H···O6 | 124.37 | 99.33 | 2.43 | | N2-H···O6 | 125.84 | 98.05 | 3.01 |
| | N2-H···N7 | 156.66 | 110.34 | 1.91 | | N2-H···N7 | 159.93 | 124.71 | 2.04 |
| **GC♦C** | N4-H···O2 | 163.55 | 107.65 | 1.97 | **GC♦C** | N4-H···O2 | 152.53 | 114.33 | 2.19 |
| | N4-H···O6 | 155.26 | 117.89 | 2.37 | | N4-H···O6 | 151.06 | 114.80 | 2.61 |
| | N4-H···N7 | 141.34 | 96.88 | 2.94 | | N4-H···N7 | 147.84 | 110.80 | 2.56 |
| **TA♦A** | N6-H···O4 | 168.47 | 142.11 | 2.18 | **TA♦A** | N6-H···O4 | 168.47 | 142.11 | 2.18 |
| | N6-H···N1 | 130.02 | 91.81 | 2.49 | | N6-H···N1 | 141.47 | 113.18 | 2.95 |
| **AT♦T** | N6-H···O4 | 124.37 | 148.73 | 2.18 | **AT♦T** | N6-H···O4 | 171.12 | 153.71 | 2.60 |
| | N3-H···O4 | 162.27 | 139.10 | 2.11 | | N3-H···O4 | 176.18 | 161.04 | 2.53 |

[a]DHA (Donor–H···Acceptor): Ideal hydrogen bond angle is close to 180°. Effective hydrogen bonds are generally bigger than 150°.

[b]HAY (H···Acceptor–Y): Orientation of lone-pair electrons on the acceptor atom. The ideal angle for both nitrogen and oxygen is 120°. Effective hydrogen bonds are generally 100–140°.

**Table S3.** Lengths of the homologous strand and the third strand of HP-triplexes.

| B-form triplex (10 nt) | | | Z-form triplex (12 nt) | | |
|---|---|---|---|---|---|
| Distance of phosphate[a] | Homologous strand (Å) | Third strand (Å) | Distance of phosphate | Homologous strand (Å) | Third strand (Å) |
| 1T-2C | 6.739 | 7.341 | 1C-2G | 6.708 | 8.009 |
| 2C-3T | 6.566 | 9.244 | 2G-3T | 7.024 | 7.734 |
| 3T-4T | 6.567 | 8.795 | 3T-4G | 7.449 | 8.749 |
| 4T-5C | 6.717 | 7.761 | 4G-5C | 7.612 | 7.586 |
| 5C-6G | 6.813 | 8.792 | 5C-6G | 5.765 | 8.261 |
| 6G-7A | 6.643 | 9.390 | 6G-7T | 7.214 | 7.035 |
| 7A-8T | 6.384 | 8.426 | 7T-8A | 6.412 | 8.147 |
| 8T-9T | 6.366 | 8.762 | 8A-9C | 8.378 | 7.048 |
| 9T-10C | 6.599 | 7.786 | 9C-10G | 6.868 | 8.818 |
| 10C-11T | 6.788 | 8.352 | 10G-11C | 8.452 | 7.501 |
| | | | 11C-12G | 6.917 | 8.722 |
| | | | 12G-13C | 7.727 | 6.681 |
| **1T-11T**[b] | 66.182 | 84.649 | **1C-13C** | 86.526 | 94.291 |
| **Average**[c] | 6.618 | 8.465 | **Average** | 7.211 | 7.858 |

[a]Distance between adjacent phosphate groups.

[b]The sequence of 3rd strand for B-form: 5′-TCTTCGATTCT-5′; for the Z-form 3rd strand: 5′-CGTGCGTACGCGCG-5′.

[c]Average length per base.

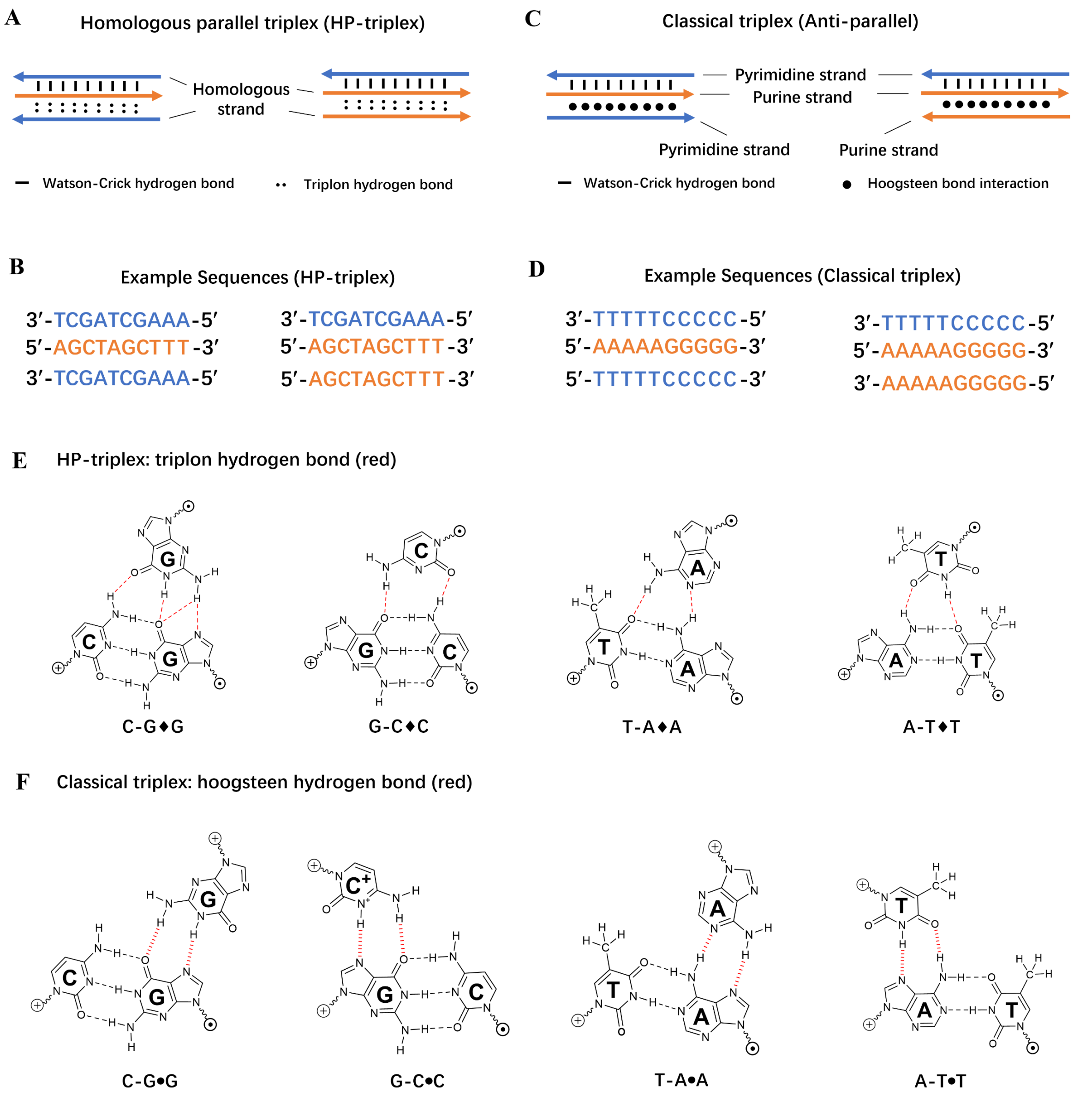


**Figure S1.** Structures of the HP-triplex and classical triplex. Sequence composition **(A)** and specific sequence examples **(B)** of the HP-triplex. Sequence composition **(C)** and specific sequence examples **(D)** of the classical triplex (anti-parallel). **(E)** Schematic of hydrogen bonding of triplons within the HP-triplex. Black lines denote Watson–Crick base pairs, and red lines represent triplon hydrogen bonds[1,2]. **(F)** Hydrogen-bonding arrangements of triplet bases in the classical triplex. Black lines denote hydrogen bonds in Watson–Crick base pairs, and red lines indicate Hoogsteen hydrogen bonds[3].

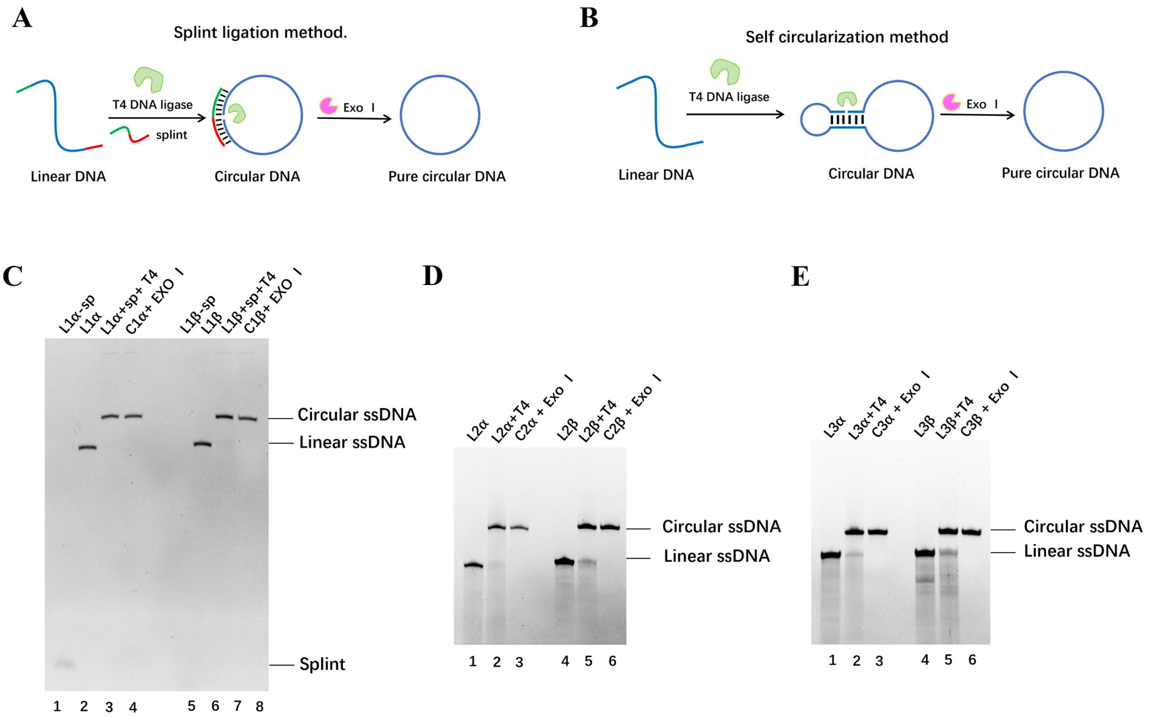


**Figure S2. Results for circularization of linear ssDNA.** (**A**) Schematic of ligation using a splint. The 3′-OH and 5′-phosphate of linear ssDNA were brought into close proximity via the complementary splint, and ligation was performed by T4 DNA ligase. Linear by-products were removed by exonuclease I digestion. **(B)** Schematic of self-circularization without a splint. The 3′-OH and 5′-phosphate of linear ssDNA were brought into close proximity through self-folding before ligation by T4 DNA ligase. Linear by-products were removed by exonuclease I digestion[4]. Results of ligation by the splint **(C)** or the self-circularization **(D, E)**. **(C)** Lanes 1, 5： splint； Lanes 2, 6： linear ssDNA； Lanes 3, 7： linear ssDNA mixed with the splint and T4 DNA ligase； Lanes 4, 8： circularized ssDNA treated with Exonuclease I. Conditions: 2 μM ssDNA, 4 μM Splint. **(D, E)** Lanes 1, 4： linear ssDNA； Lanes 2, 5： linear ssDNA mixed with T4 DNA ligase； Lanes 3, 6： circularized ssDNA treated with Exonuclease I. Conditions: 10 μM ssDNA; 15% dPAGE containing 8.0 M urea.

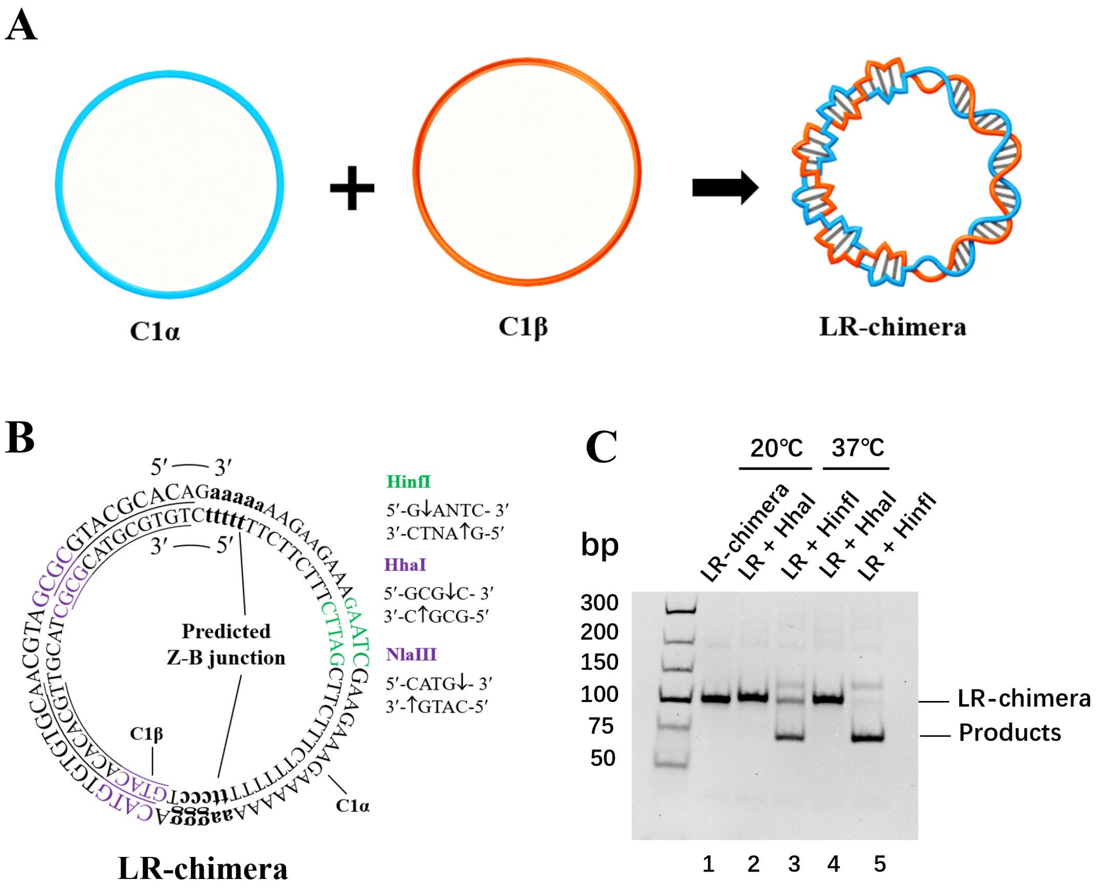


**Figure S3. Validation of the LR-chimera formation by restriction endonuclease digestion. (A)** Scheme for LR-chimera formation. C1α and C1β were hybridized at a molar ratio of 1:1 to form the LR-chimera. **(B)** Sequence of the LR-chimera and its restriction enzyme sites. Underlined regions: APP sequences (forming Z-DNA); green: HinfI restriction site; purple: HhaI and NlaIII restriction sites; lowercase letters: predicted Z–B junction sites (typically 1 bp). **(C)** Digestion of LR-chimera by HhaI or HinfI. Lane 1: LR-chimera; Lanes 2, 4: LR-chimera incubated with HhaI; Lanes 3, 5: LR-chimera incubated with HinfI. Incubation was performed at 20 °C for 1 h or at 37 °C for 10 min. Other conditions: 0.1 μM DNA, 0.5 U/μL restriction enzyme; 10% native-PAGE (10 mM $MgCl_2$, pH 7.5), run at 10 °C, stained with GelRed.

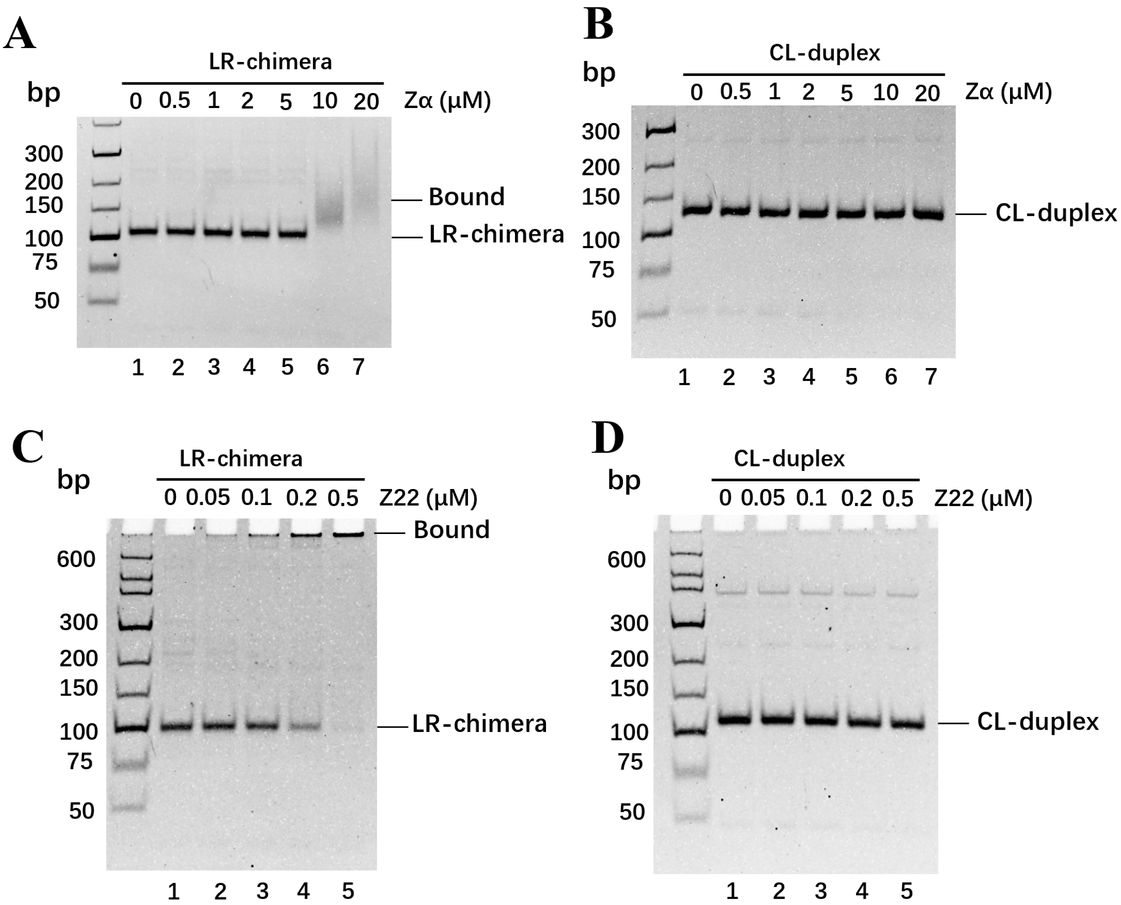


**Figure S4. Validation of Z-DNA structure within the LR-chimera using Zα and Z22.** Binding results of Zα to the LR-chimera **(A)** and CL-duplex (nicked circular duplex with identical sequence) **(B)**. Lane 1: dsDNA (LR-chimera or CL-duplex); Lanes 2–7: dsDNA incubated with Zα at varying concentrations (0–20 μM). Binding results of Z22 to the LR-chimera **(C)** and CL-duplex **(D)**. Lane 1: dsDNA (LR-chimera or CL-duplex); Lanes 2–7: dsDNA incubated with Z22 at varying concentrations (0–0.5 μM). Conditions: 10 mM $MgCl_2$, 10 mM HEPES buffer (pH 7.0), 20 °C, 2 h; 10% native-PAGE (10 mM $MgCl_2$, pH 7.5) at 10 °C, stained with GelRed.

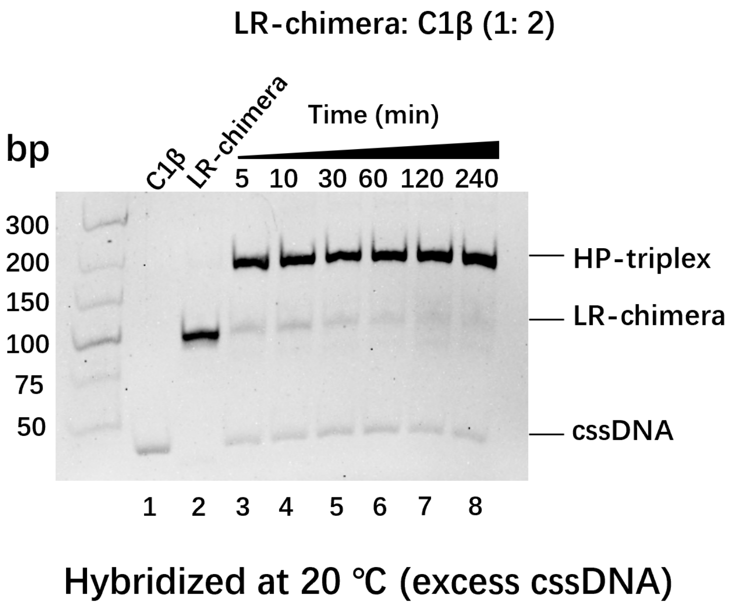


**Figure S5. Electrophoresis analysis of hybridization between LR-chimera and 2-fold excess of C1β.** Lane 1: C1β (homologous cssDNA); Lane 2: LR-chimera; Lanes 3–8: LR-chimera incubated with C1β at a 1:2 molar ratio at 20 °C for various durations (5–240 min). Other conditions: 0.1 μM LR-chimera, 0.2 μM C1β, 10 mM $MgCl_2$, 10 mM HEPES (pH 7.0); 10% native-PAGE (10 mM $MgCl_2$, pH 7.5) at 10 °C, stained with GelRed.

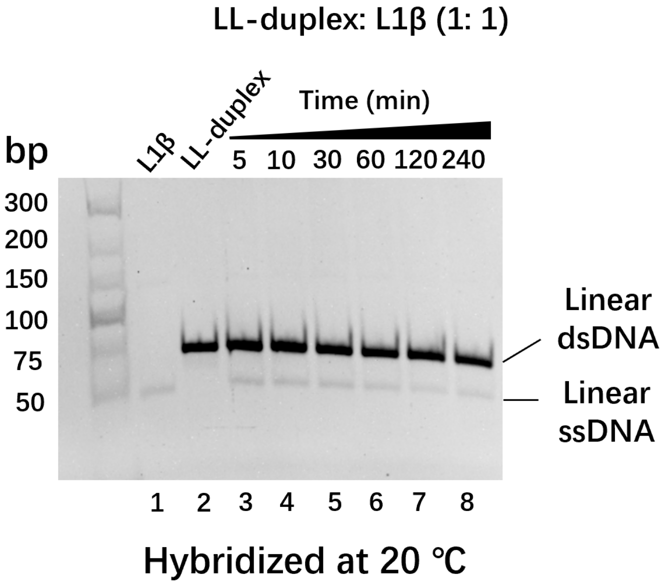


**Figure S6. Results for mixtures of linear ssDNA and linear duplex (LL-duplex).** Lane 1: L1β (homologous linear ssDNA); Lane 2: LL-duplex; Lanes 3–8: LL-duplex incubated with L1β at a 1:1 molar ratio at 20 °C for different durations (5–240 min). Conditions: 0.1 μM DNA; 10 mM $MgCl_2$, 10 mM HEPES (pH 7.0). 10% native-PAGE (10 mM $MgCl_2$, pH 7.5) at 10 °C, stained with GelRed.

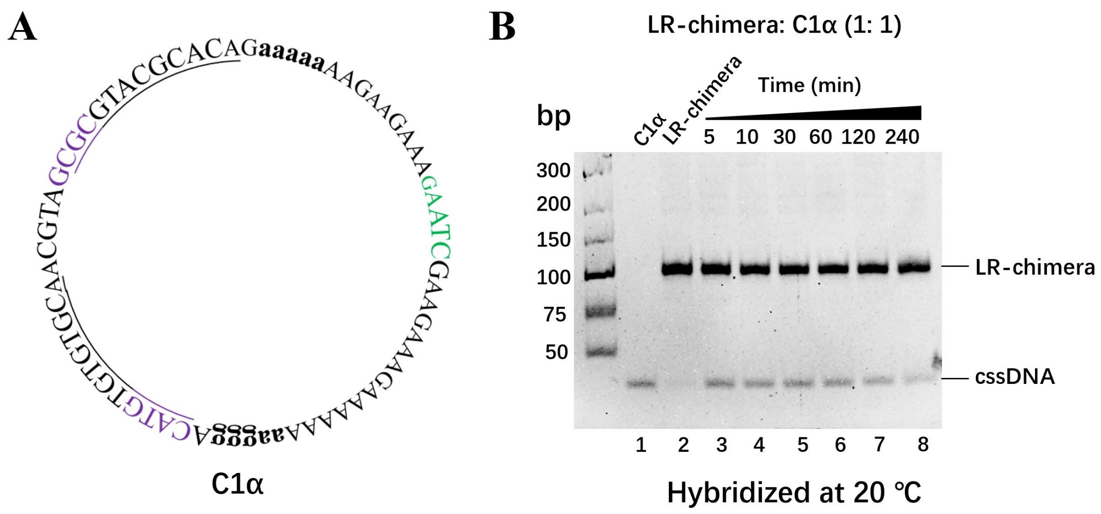


**Figure S7. Results for mixtures of LR-chimera and C1α. (A)** Schematic diagram of C1α (purine-rich homologous cssDNA). **(B)** Lane 1: C1α; Lane 2: LR-chimera; Lanes 3–8: C1α incubated with LR-chimera (1:1) at 20 °C for different durations (5–240 min). Conditions: 0.1 µM DNA; 10 mM $MgCl_2$, 10 mM HEPES (pH 7.0). 10% native-PAGE (10 mM $MgCl_2$, pH 7.5) at 10 °C, stained with GelRed.

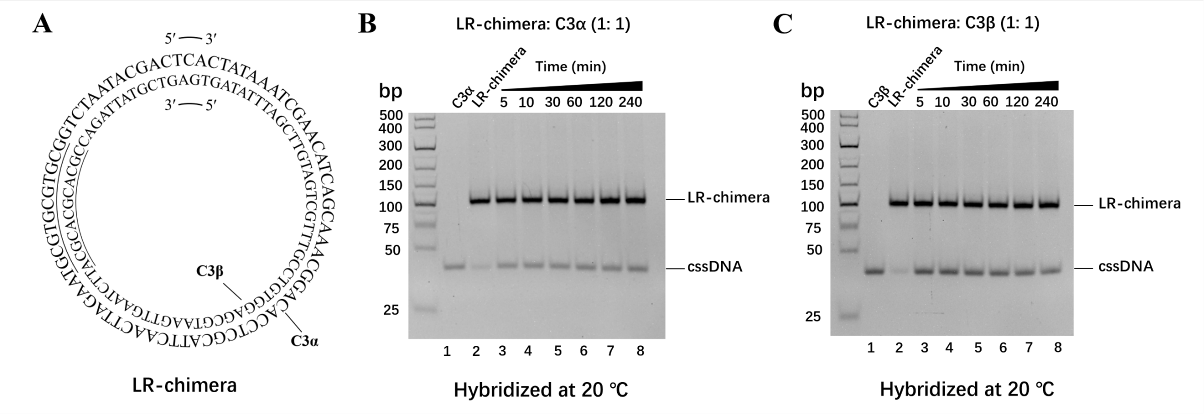


**Figure S8. Results for mixtures of LR-chimera and homologous cssDNAs with random sequences. (A)** Schematic of the random sequence. Underlined regions: APP sequences; the right segment corresponds to a random sequence with relatively even distribution of the four bases. This sequence was obtained from our previous work and designated cc74[5]. Results for incubation of LR-chimera with C3α **(B)** or C3β **(C)**. Lane 1: homologous cssDNA; Lane 2: LR-chimera; Lanes 3–8: cssDNA incubated with LR-chimera (1:1) at 20 °C for different durations (5–240 min). Conditions: 0.1 μM DNA; 10 mM $MgCl_2$, 10 mM HEPES (pH 7.0). 10% native-PAGE (10 mM $MgCl_2$, pH 7.5) at 10 °C, stained with GelRed.

Zhong et al.

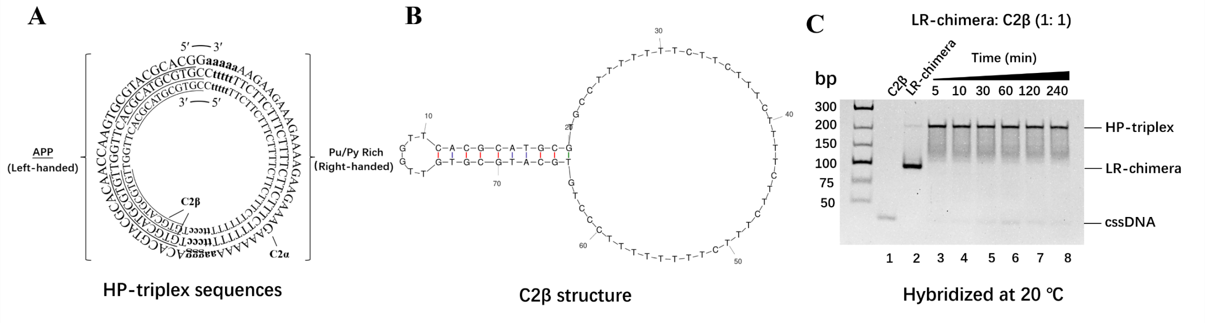


**Figure S9. Results for mixtures of LR-chimera and homologous cssDNA bearing a hairpin on the left side.** **(A)** Schematic of the cssDNA sequence that forms the HP-triplex. Underlined regions: APP sequences; the right segment: poly-Pu/Py sequence. Lowercase bases: predicted range for Z–B junction formation. **(B)** Schematic of C2β (homologous cssDNA). The Z-DNA-forming region contains a 9 bp hairpin structure. **(C)** Hybridization results for C2β with LR-chimera. Lane 1: C2β; Lane 2: LR-chimera; Lanes 3–8: C2β incubated with LR-chimera (1:1) at 20 °C for different durations (5–240 min). Conditions: 0.1 μM DNA; 10 mM $MgCl_2$, 10 mM HEPES (pH 7.0). 10% native-PAGE (10 mM $MgCl_2$, pH 7.5) at 10 °C, stained with GelRed.

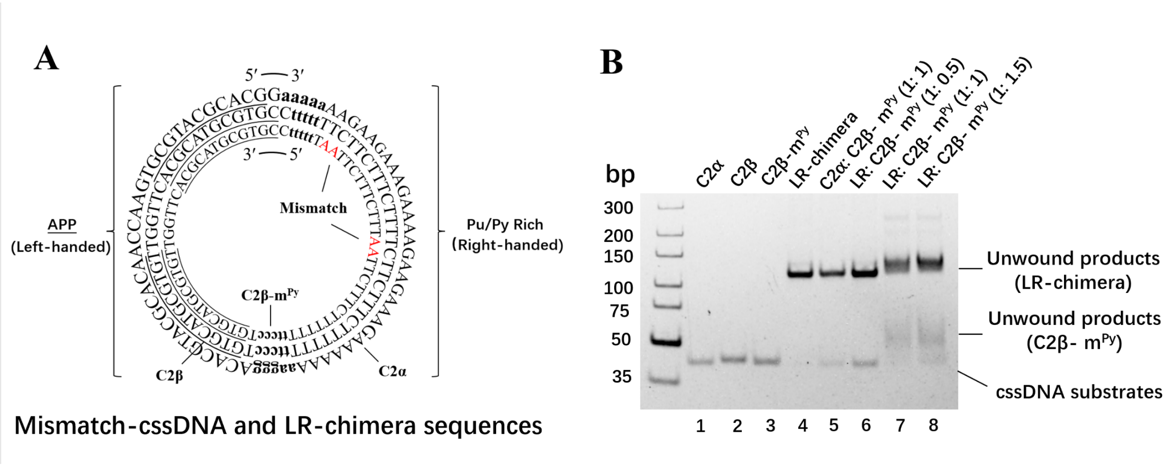


**Figure S10. Effects of mismatched cssDNA on HP-triplex formation.** (A) Schematic of C2β-m$^{Py}$ and LR-chimera. Underlined regions: APP sequences; the right segment: poly-PuPy sequence; red AA bases: mismatch sites of C2β-m$^{Py}$. (B) Hybridization results for C2β-m$^{Py}$ with LR-chimera. Lanes 1, 2: C2α and C2β; Lane 3: C2β-m$^{Py}$; Lane 4: LR-chimera; Lane 5: C2β-m$^{Py}$ incubated with C2α (1:1); Lanes 6–8: LR-chimera incubated with C2β-m$^{Py}$ (0.5-, 1-, and 1.5-fold molar excess) at 20 °C for 4 h. Conditions: 0.1 µM DNA; 10 mM $MgCl_2$, 10 mM HEPES (pH 7.0). 10% native-PAGE (10 mM $MgCl_2$, pH 7.5) at 10 °C, stained with GelRed.

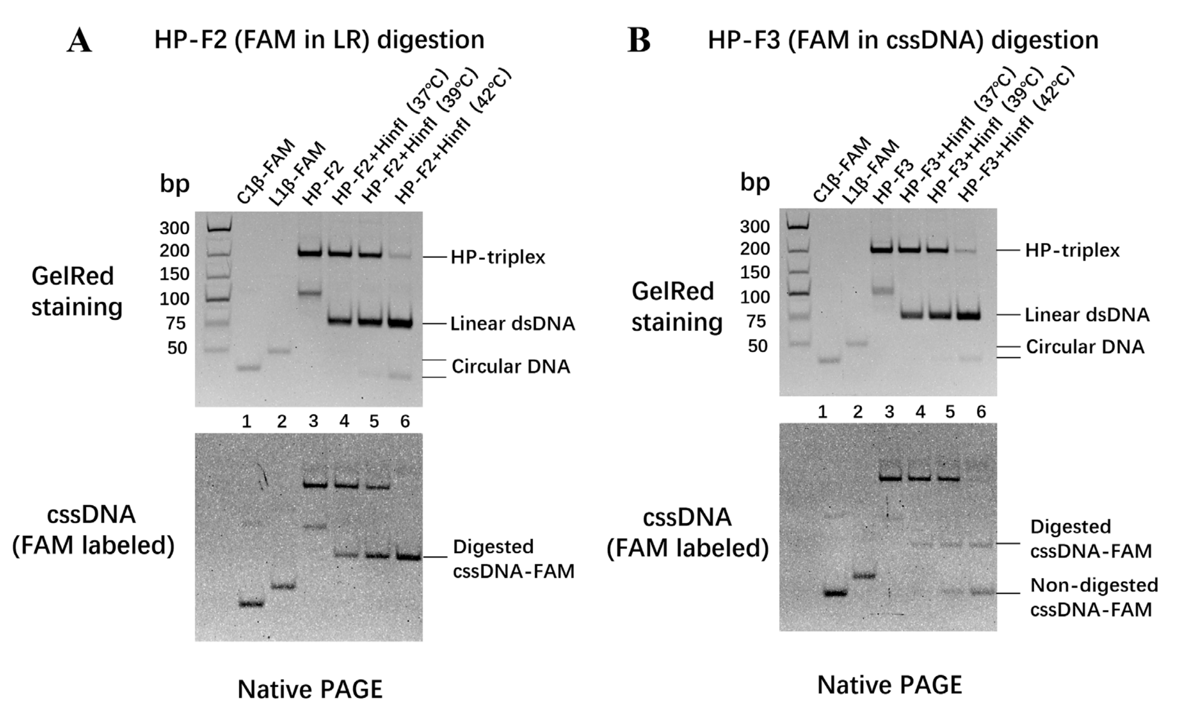


**Figure S11. Analysis of high-temperature digestion products of HP-triplex.** Digestion results for HP-F2 **(A)** and HP-F3 **(B)** at different temperatures. Lane 1: C1β-FAM; Lane 2: L1β-FAM; Lane 3: FAM-labeled HP-triplex (HP-F2 or HP-F3); Lanes 4–6: FAM-labeled HP-triplex digested at 37 °C, 39 °C, or 42 °C. Lower panel: Direct FAM fluorescence imaging for tracking FAM-labeled cssDNA. Upper panel: GelRed staining of the same gel shown in the lower panel. Conditions: 0.1 µM DNA, 0.5 U/µL restriction enzyme, incubation at 37–42 °C for 12 h. 10% native-PAGE (10 mM $MgCl_2$, pH 7.5) at 10 °C.

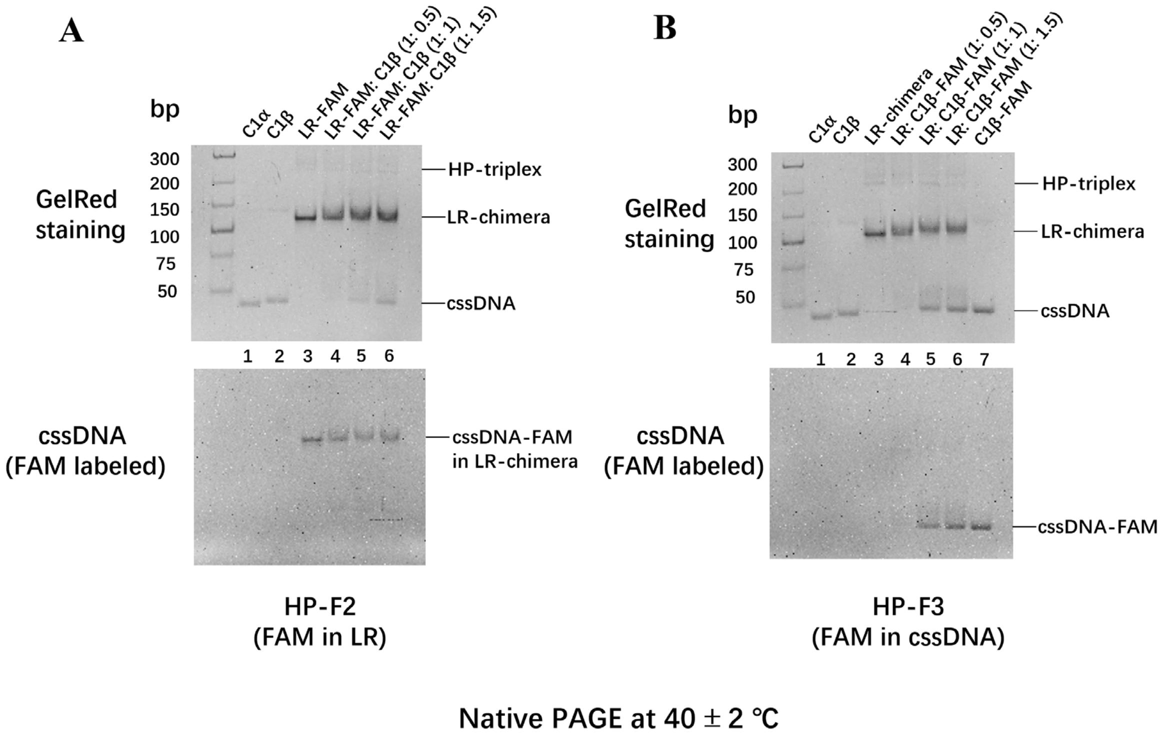


**Figure S12. Electrophoresis results for the HP-triplex at 40 °C.** Electrophoresis of HP-F2 **(A)** and HP-F3 **(B)**. Lanes 1 and 2: C1α, C1β; Lane 3: LR-chimera-FAM; Lanes 4–6: LR-chimera-FAM mixed with C1β (panel A) and LR-chimera mixed with C1β-FAM (panel B) at molar ratios of 0.5, 1, and 1.5; Lane 7: C1β-FAM. Lower panel: FAM fluorescence imaging; Upper panel: GelRed-stained image of the same gel shown in the lower panel. Conditions: 0.1 μM DNA; 10% native-PAGE (10 mM $MgCl_2$, pH7.5) at 40°C.

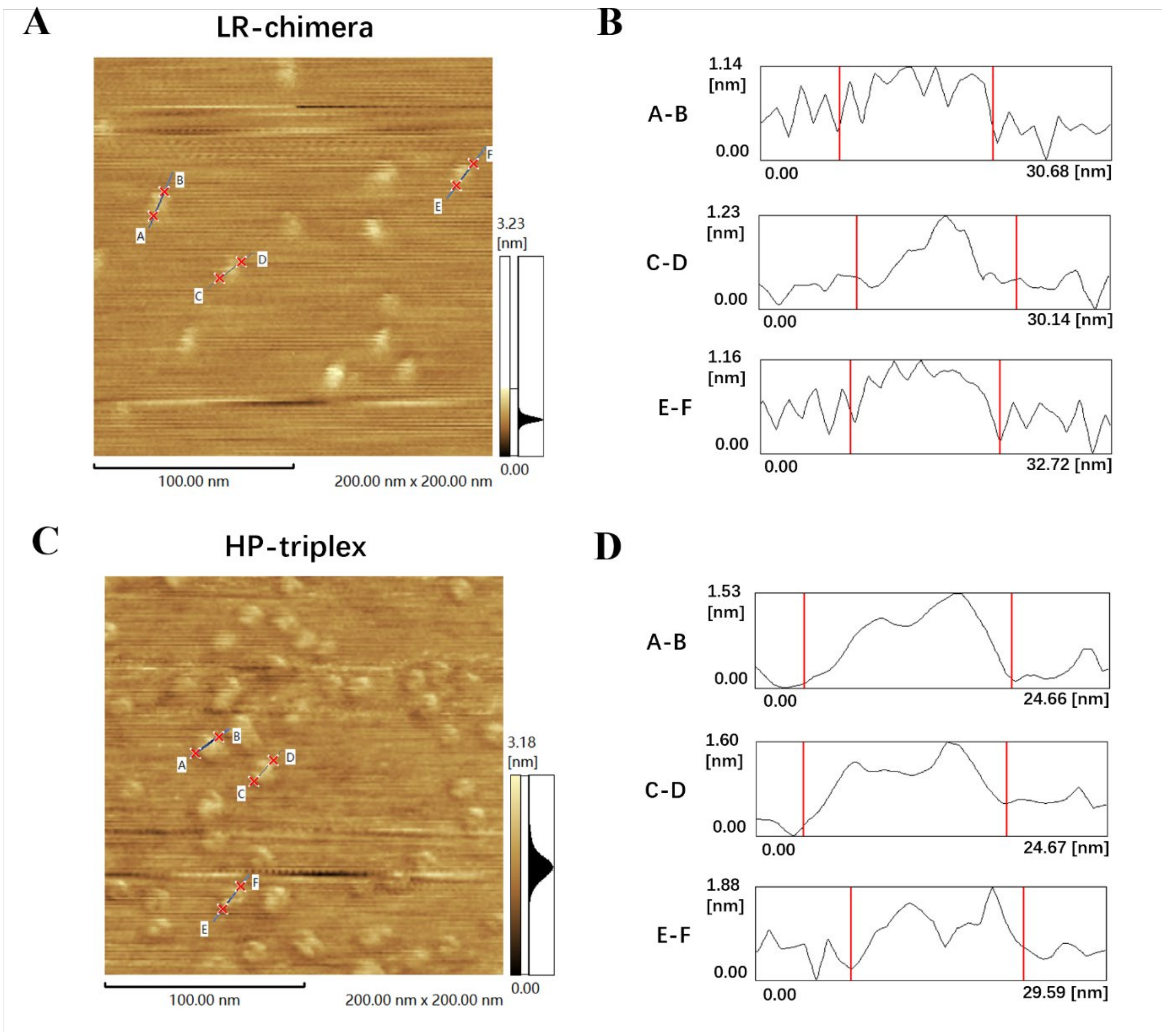


**Figure S13. AFM imaging of DNA structures. (A)** AFM image of the LR-chimera. **(B)** Height statistical analysis of the LR-chimera shown in (A). **(C)** AFM image of the HP-triplex. **(D)** Height statistical analysis of the HP-triplex shown in (C). 10 nM DNA; scale bar: 100 nm.

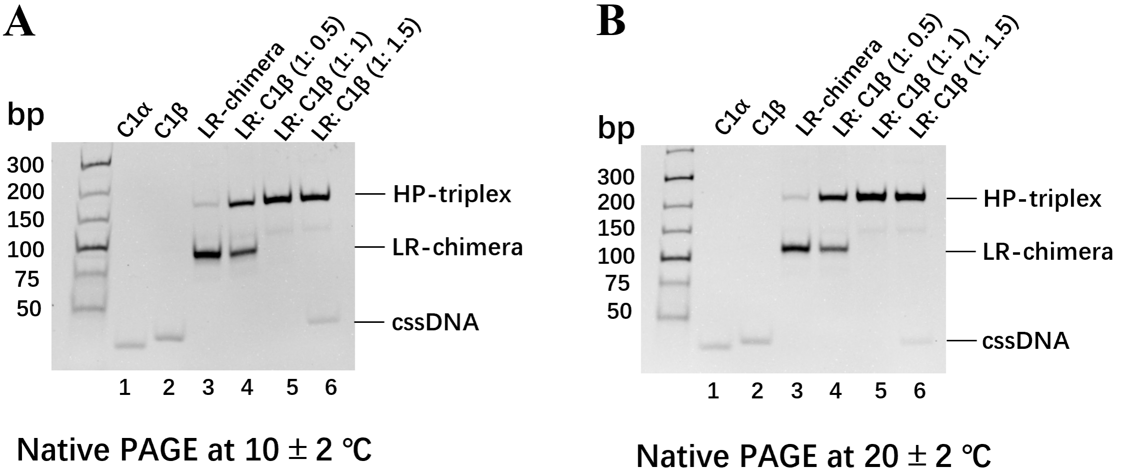


**Figure S14. Stability analysis of the HP-triplex using 10% native-PAGE.** Electrophoresis results for the HP-triplex at 10 °C **(A)** and 20 °C **(B)**. Lanes 1, 2: C1α and C1β; Lane 3: LR-chimera; Lanes 4–6: LR-chimera incubated with C1β (0.5-, 1-, and 1.5-fold). Conditions: 0.1 μM DNA; 10 mM $MgCl_2$, 10 mM HEPES (pH 7.0). Gels were stained with GelRed.

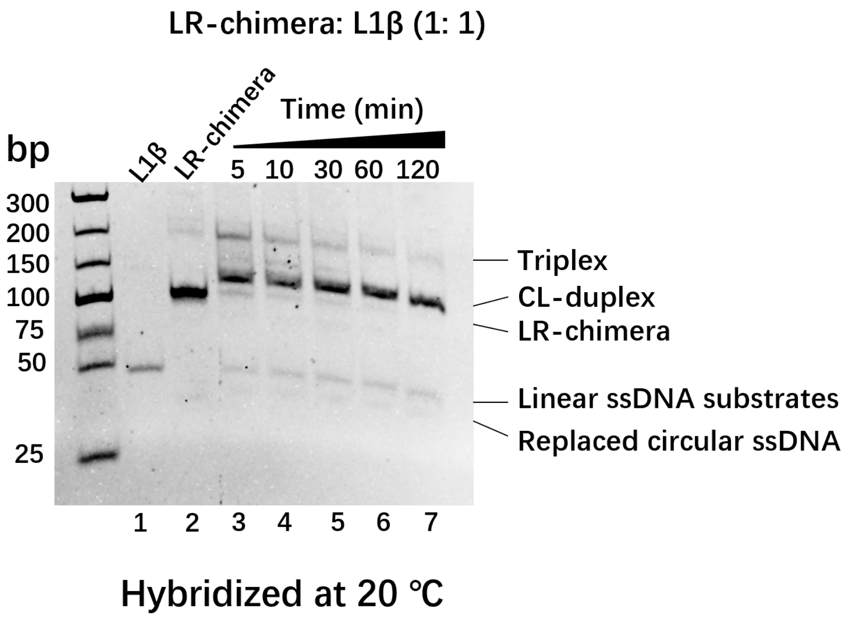


**Figure S15. Hybridization results for LR-chimera with L1β.** Lane 1: L1β (homologous linear ssDNA); Lane 2: LR-chimera; Lanes 3–7: LR-chimera incubated with L1β (1:1) at 20 °C for different durations (5–120 min). Conditions: 0.1 μM DNA; 10 mM $MgCl_2$, 10 mM HEPES (pH 7.0). 10% native-PAGE (10 mM $MgCl_2$, pH 7.5) at 10 °C, stained with GelRed.

Zhong et al.

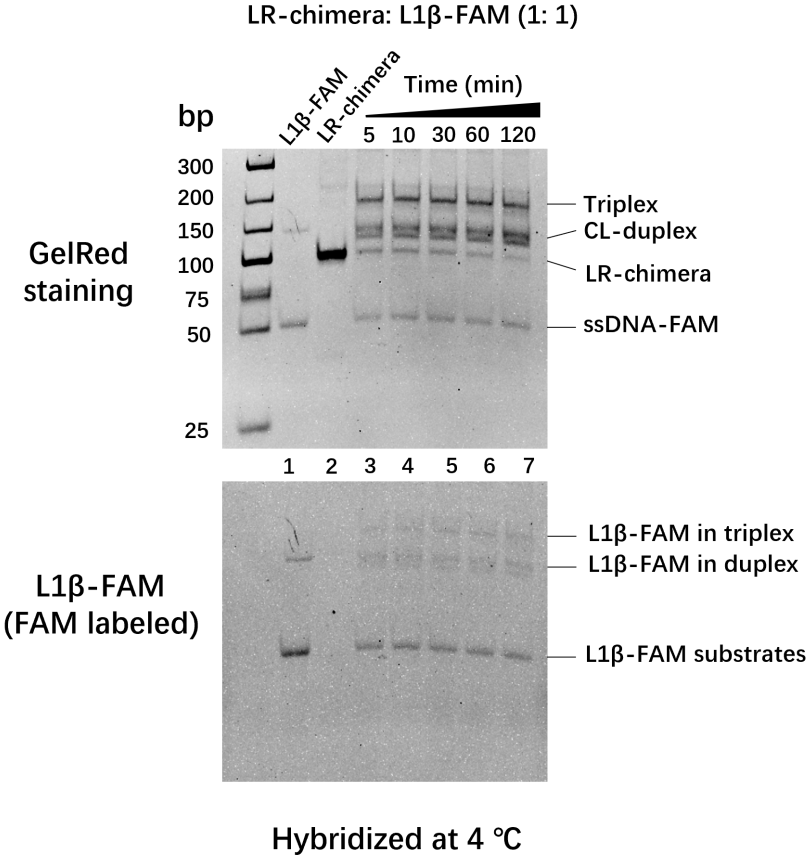


**Figure S16. Hybridization results for LR-chimera with L1β-FAM.** Lane 1: L1β-FAM (FAM-labeled linear ssDNA); Lane 2: LR-chimera; Lanes 3–7: LR-chimera incubated with L1β-FAM (1:1) at 4 °C for different durations (5–120 min). Lower panel: FAM fluorescence imaged; Upper panel: the same gel shown in the lower panel stained with GelRed. Conditions: 0.1 μM DNA; 10 mM $MgCl_2$, 10 mM HEPES (pH 7.0). 10% native-PAGE (10 mM $MgCl_2$, pH 7.5) at 4 °C.

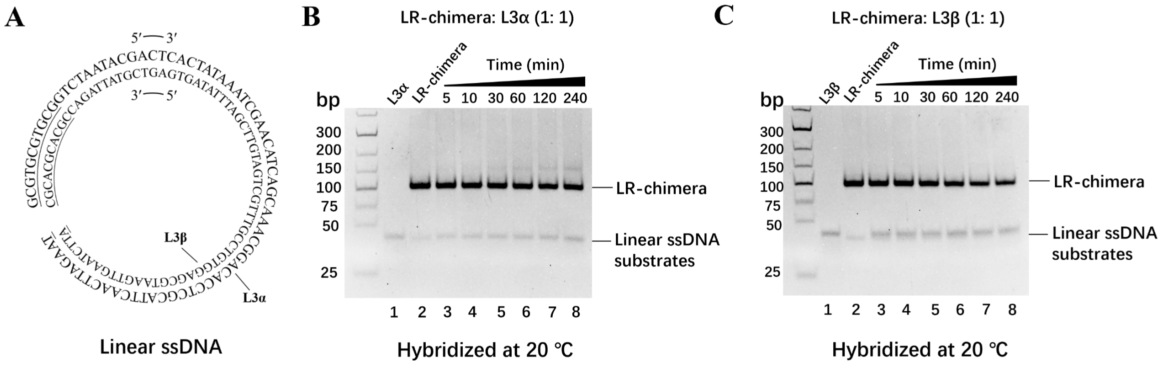


**Figure S17. Results for mixtures of LR-chimera and linear ssDNA with random sequences. (A)** Schematic of two complementary linear DNA strands (L3α and L3β). Results for incubation of LR-chimera with L3α **(B)** or L3β **(C)**. Lane 1: linear ssDNA (L3α or L3β); Lane 2: LR-chimera; Lanes 3–7: LR-chimera incubated with linear ssDNA (1:1) at 20 °C for different durations (5–120 min). Conditions: 0.1 μM DNA; 10 mM $MgCl_2$, 10 mM HEPES (pH 7.0). 10% native-PAGE (10 mM $MgCl_2$, pH 7.5) at 10 °C, stained with GelRed.

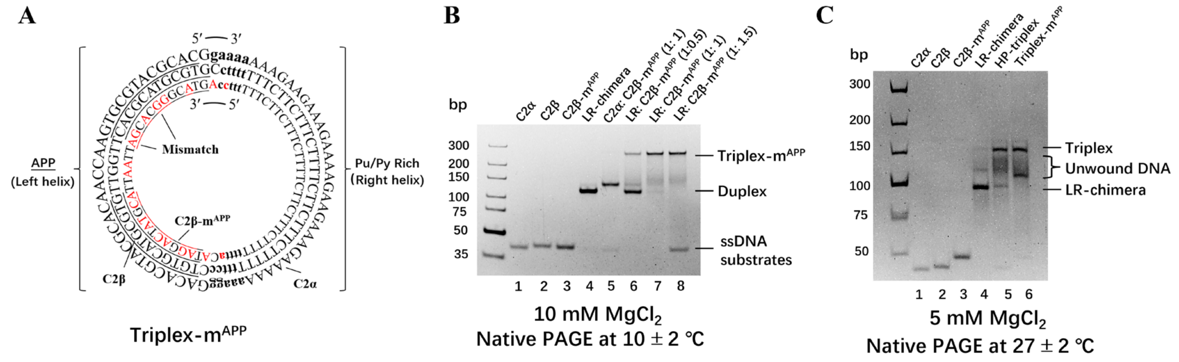


**Figure S18. Effects of mismatch on the Z-DNA-side of 3rd strand within the HP-triplex. (A)** Schematic of mismatch sequences in 3rd strand. C2β-m$^{APP}$ contains 20 bp mismatches on the APP side relative to C2β. Red labels indicate mismatch sites. **(B)** Results for mixtures of C2β-m$^{APP}$ and LR-chimera. Lanes 1–3: C2α, C2β, C2β-m$^{APP}$; Lane 4: LR-chimera; Lane 5: C2β-m$^{APP}$ incubated with C2α (1:1); Lanes 6–8: LR-chimera incubated with C2β-m$^{APP}$ (0.5-, 1-, and 1.5-fold). Conditions: 0.1 µM DNA; buffer: 10 mM HEPES (10 mM $MgCl_2$, pH 7.0). 10% native-PAGE (10 mM $MgCl_2$, pH 7.5) at 10 °C. **(C)** Electrophoretic results for Triplex-m$^{APP}$ and HP-triplex at 27 °C. Lanes 1–3: C2α, C2β, C2β-m$^{APP}$; Lane 4: LR-chimera; Lane 5: HP-triplex; Lane 6: Triplex-m$^{APP}$. Conditions: 0.1 µM DNA; 10 mM HEPES (5 mM $MgCl_2$, pH 7.0). 10% native-PAGE (5 mM $MgCl_2$, pH 7.5) at 27 ± 2 °C, stained with GelRed.

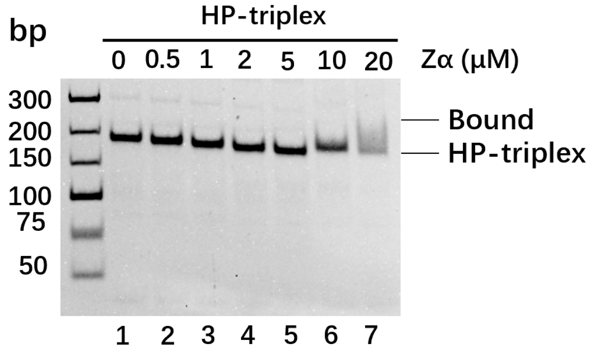


**Figure S19. Binding results for Zα to the HP-triplex assembled via the C1 circle.** Lane 1: HP-triplex; Lanes 2–7: HP-triplex (0.1 μM) incubated with Zα at varying concentrations (0.5–20 μM) at 20 °C for 2 h. 10 mM HEPES (10 mM $MgCl_2$, pH 7.0). 10% native-PAGE (10 mM $MgCl_2$, pH 7.5) at 10 °C, stained with GelRed.

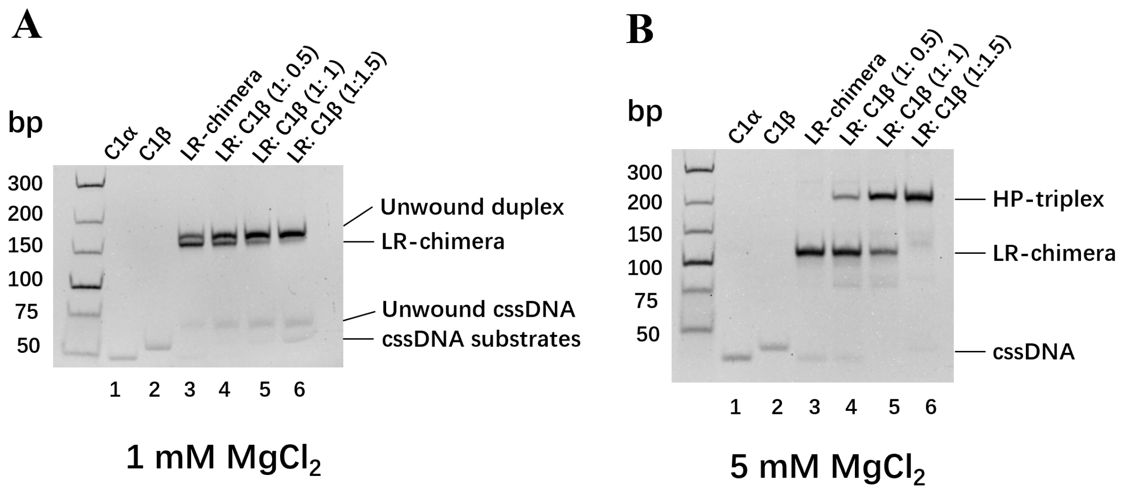


**Figure S20. Effects of magnesium ion concentrations on the HP-triplex structure**. PAGE detection results at 1 mM $MgCl_2$ **(A)** and 5 mM $MgCl_2$ **(B).** Lanes 1, 2: C1α, C1β; Lane 3: LR-chimera; Lanes 4–6: LR-chimera incubated with C1β (0.5-, 1-, and 1.5-fold) at 20 °C for 4 h. Conditions: 0.1 μM DNA; 10 mM HEPES (1 mM $MgCl_2$ or 5 mM $MgCl_2$, pH 7.0); 10% native-PAGE (1 mM $MgCl_2$ or 5 mM $MgCl_2$, pH 7.5) at 10 °C, stained with GelRed.

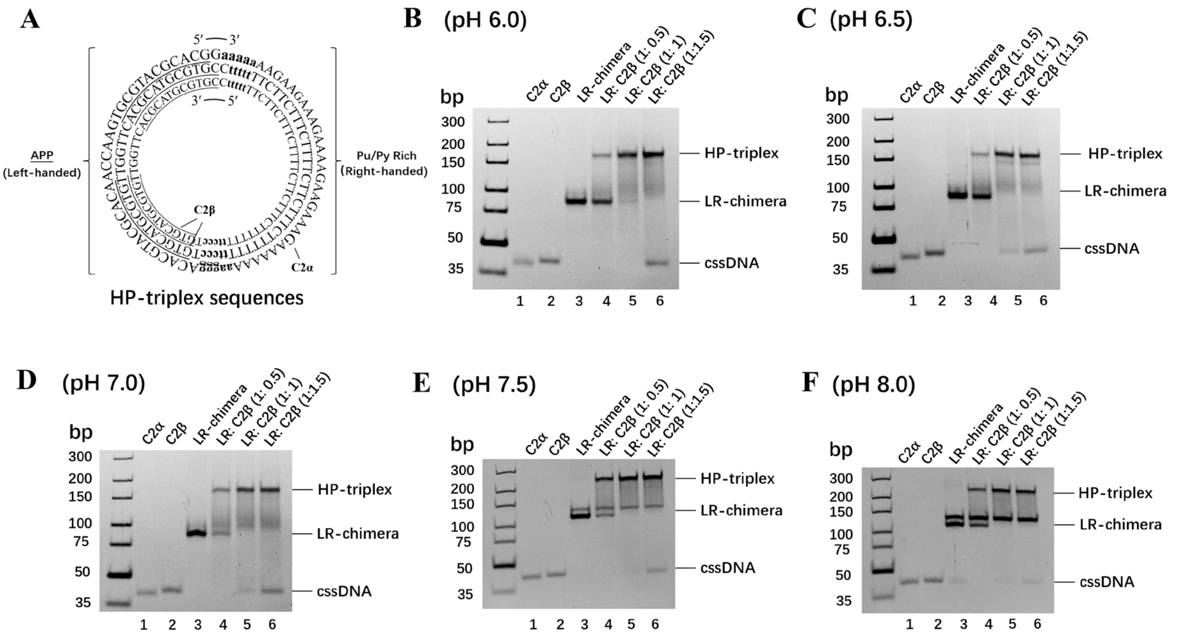


**Figure S21. Effects of pH (6.0–8.0) on the HP-triplex structure. (A)** Sequence of HP-triplex. **(B–F)** 10% native-PAGE analysis of HP-triplex formation at pH values ranging from 6.0 to 8.0. Lanes 1, 2: C2α, C2β; Lane 3: LR-chimera; Lanes 4–6: LR-chimera incubated with C2β (0.5-, 1-, and 1.5-fold) at 20 °C for 4 h. 0.1 μM DNA; 10 mM $MgCl_2$, 10 mM HEPES (pH 6.0–8.0); 10% native-PAGE (10 mM $MgCl_2$, pH 7.5) at 10 °C, stained with GelRed.

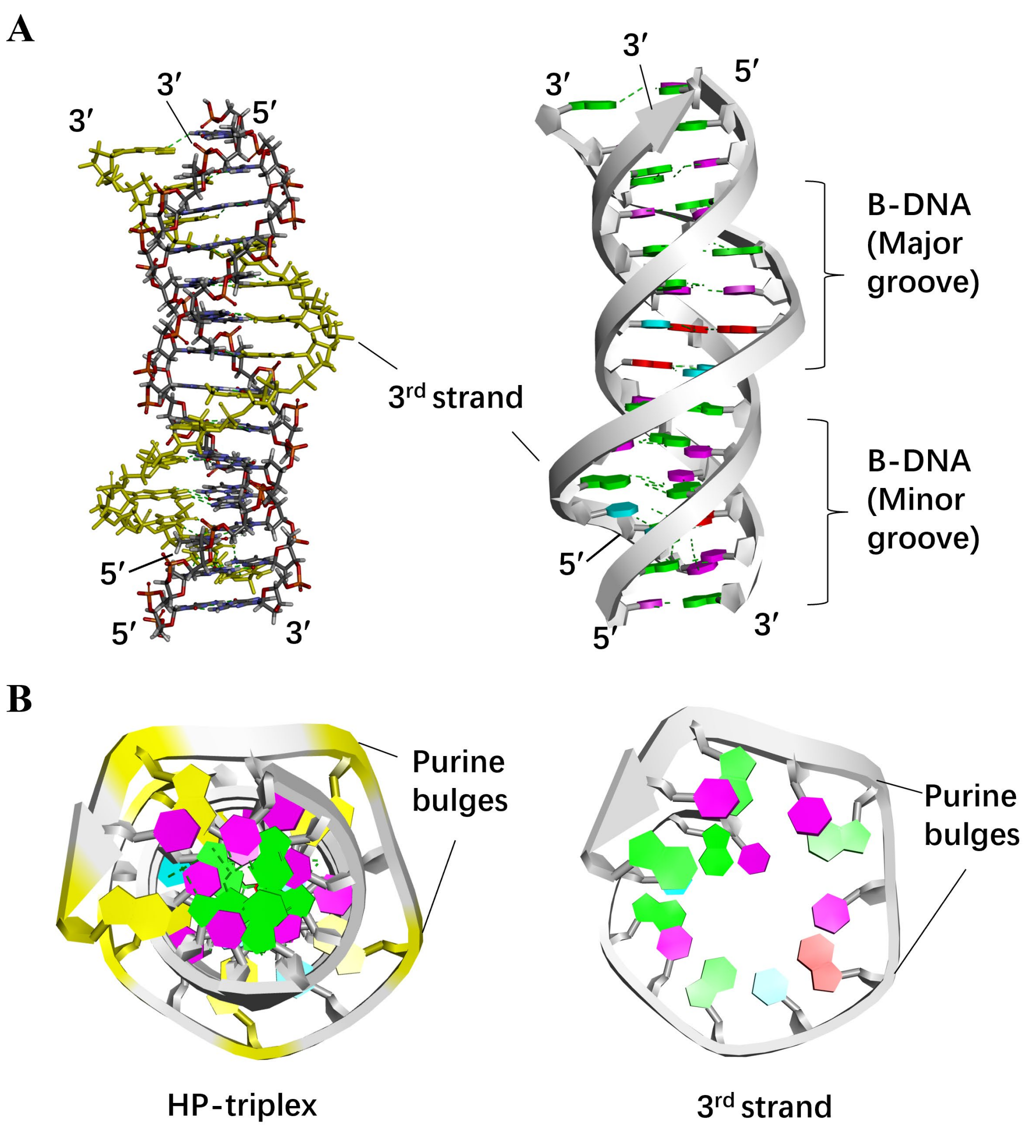


**Figure S22. Molecular model of the right-handed HP-triplex formed by the APP sequence. (A)** Structural model of the triplex. Left: stick view; Right: ribbon view. The sequence of the homologous third strand is 5′-CGTGCGTACGCGCG-3′. The 3rd strand resides within the major groove of the duplex. Bases are shown in different colors: red, A; light blue, T; purple, C; green, G. **(B)** Top view of panel A. Left: overall triplex view, with purine bases in the 3rd strand highlighted in yellow. Right: view along 3rd strand. Purine bases are located at the bulge sites.

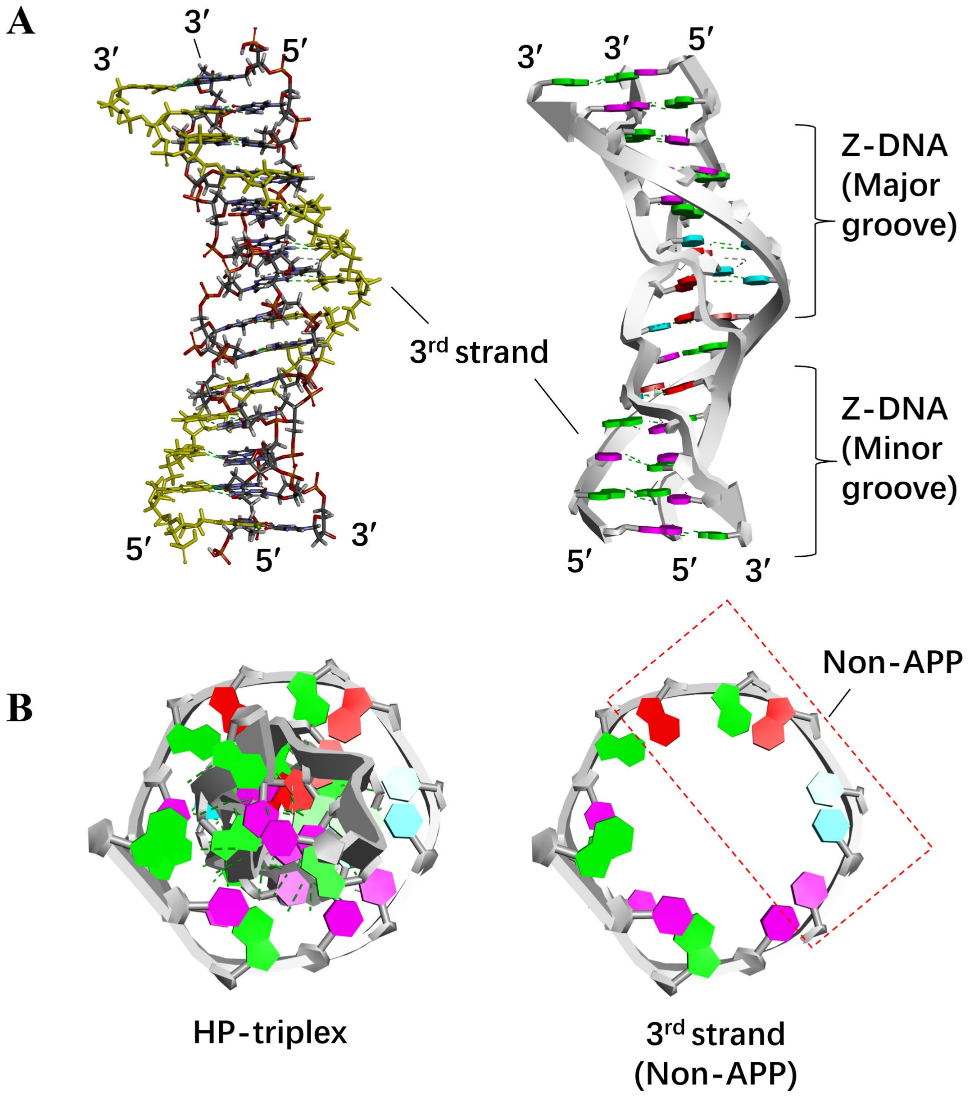


**Figure S23. Molecular model of the left-handed HP-triplex formed by the non-APP sequence. (A)** Structural model of the triplex. Left: stick view; Right: ribbon view. The sequence of the homologous 3rd strand is 5′-CGCGAGATTCCGCG-3′, with a central 6 bp segment (AGATTC) corresponding to the non-APP sequence. Bases are rendered in different colors: red, A; light blue, T; purple, C; green, G. **(B)** Top view of panel A. Left: overall triplex view. Right: view along 3rd strand. The non-APP sequence region is outlined by a red dashed.
